# Naïve arthritogenic SKG T cells have a defect in anergy and a repertoire pruned by superantigen

**DOI:** 10.1101/2022.01.13.476250

**Authors:** Judith Ashouri, Elizabeth McCarthy, Steven Yu, Noah Perlmutter, Charles Lin, Joe DeRisi, Chun Jimmie Ye, Arthur Weiss

**Author notes:** Contributed equally (Judith Ashouri, Elizabeth McCarthy).

## Abstract

How autoreactive CD4 T cells develop to cause rheumatoid arthritis remains unknown. We used a reporter for antigen-receptor signaling in the SKG autoimmune arthritis model to profile a T cell subpopulation enriched for arthritogenic naïve CD4 T cells before arthritis onset by bulk and single cell RNA and T cell antigen-receptor (TCR) sequencing. Our analyses reveal that despite their impaired proximal TCR signaling, a subset of SKG naïve CD4 T cells that have recently encountered endogenous antigen upregulate gene programs associated with positive regulation of T cell activation and cytokine signaling at higher levels than wild type cells in the pre-disease state. These arthritogenic cells also induce genes associated with negative regulation of T cell activation but do so less efficiently than wild type cells. Furthermore, their TCR sequences exhibit a previously unrecognized biased peripheral TCR Vβ repertoire likely driven by endogenous viral superantigens. These particular Vβs, known to recognize endogenous mouse mammary tumor virus (MMTV) superantigen, are further expanded in arthritic joints. Our results demonstrate that autoreactive naïve CD4 T cells which recognize endogenous viral superantigens are poised to cause disease by their altered transcriptome.

**Summary blurb:** Self-reactive SKG T cells that escaped negative selection harbor an independent defect in anergy that, together with chronic antigen stimulation, sets the stage for disease. Moreover, we propose a novel role for endogenous mouse mammary tumor virus (MMTV) superantigen in promoting arthritogenic T cell responses.

## Introduction

It is widely accepted that activation of conventional CD4 T cells that recognize specific self-antigen(s) via their TCRs is necessary for rheumatoid arthritis (RA) onset (1, 2). CD4 T cells from patients with RA differentiate into pathogenic effector cells despite their impaired TCR signaling (3–9). Yet, how these T cells subvert tolerance to cause disease remains incompletely understood. The SKG mouse model of arthritis represents a powerful tool to define this mechanism. Due to a hypomorphic mutation in ZAP70, a cytoplasmic tyrosine kinase critical for proximal TCR signaling, SKG mice exhibit impaired thymocyte negative selection resulting in a break in central tolerance and escape of autoreactive and arthritogenic CD4 T cells into the periphery (10–13). In response to an innate immune stimulus, dormant arthritogenic CD4 T cells become activated and SKG mice on the Balb/c genetic background develop an erosive inflammatory arthritis that resembles RA (10, 14). SKG CD4 T cells are sufficient and necessary to cause arthritis (10), and we have shown that adoptive transfer even of naïve SKG CD4 T cells (into immunodeficient hosts) are sufficient to trigger disease (11).

However, as in human RA, it is unclear how T cells can differentiate into pathogenic effector cells and produce frank disease in SKG mice despite severely impaired TCR signaling (10, 11, 15). While the SKG mice have a known defect in central tolerance, it is less clear whether they have an independent defect in peripheral tolerance. It is also unknown how escape of arthritogenic T cells into the periphery interacts with defective TCR signaling to produce frank autoimmune disease.

To address these questions, we previously developed the SKGNur mouse which combines the SKG model with a reporter of antigen receptor signaling, Nur77/*Nr4a1*-eGFP, that tethers GFP expression to the regulatory region of *Nr4a1* (encoding the orphan nuclear hormone receptor Nur77). Because NR4A1 is rapidly and selectively upregulated in response to antigen, but not inflammatory stimuli in T cells (16, 17), the SKGNur mouse allows us to identify antigen-reactive

T cells both before and during disease. We previously demonstrated that high expression of the Nur77-eGFP reporter in SKGNur mice identifies self-reactive naive CD4 T cells with greater arthritogenic potential before disease onset (11). We showed that these naïve CD4 T cells in SKG mice play a pathogenic role in arthritis development, in part, because of abnormally heightened responses to IL-6 (11) (**Supplementary Fig. 1A**). We proposed that chronic antigen stimulation and impaired TCR signaling interact uniquely in peripheral CD4 SKG T cells and result in lower expression of suppressor of cytokine signaling 3 (*Socs3*)—a key negative regulator of IL-6 signaling, and showed that this mechanism may operate in patients with RA (11). This led us to hypothesize that CD4 SKG T cells might exhibit dysregulated expression of a broader program of negative immune regulators leading, ultimately, to a breach in peripheral tolerance.

**Figure 1.**
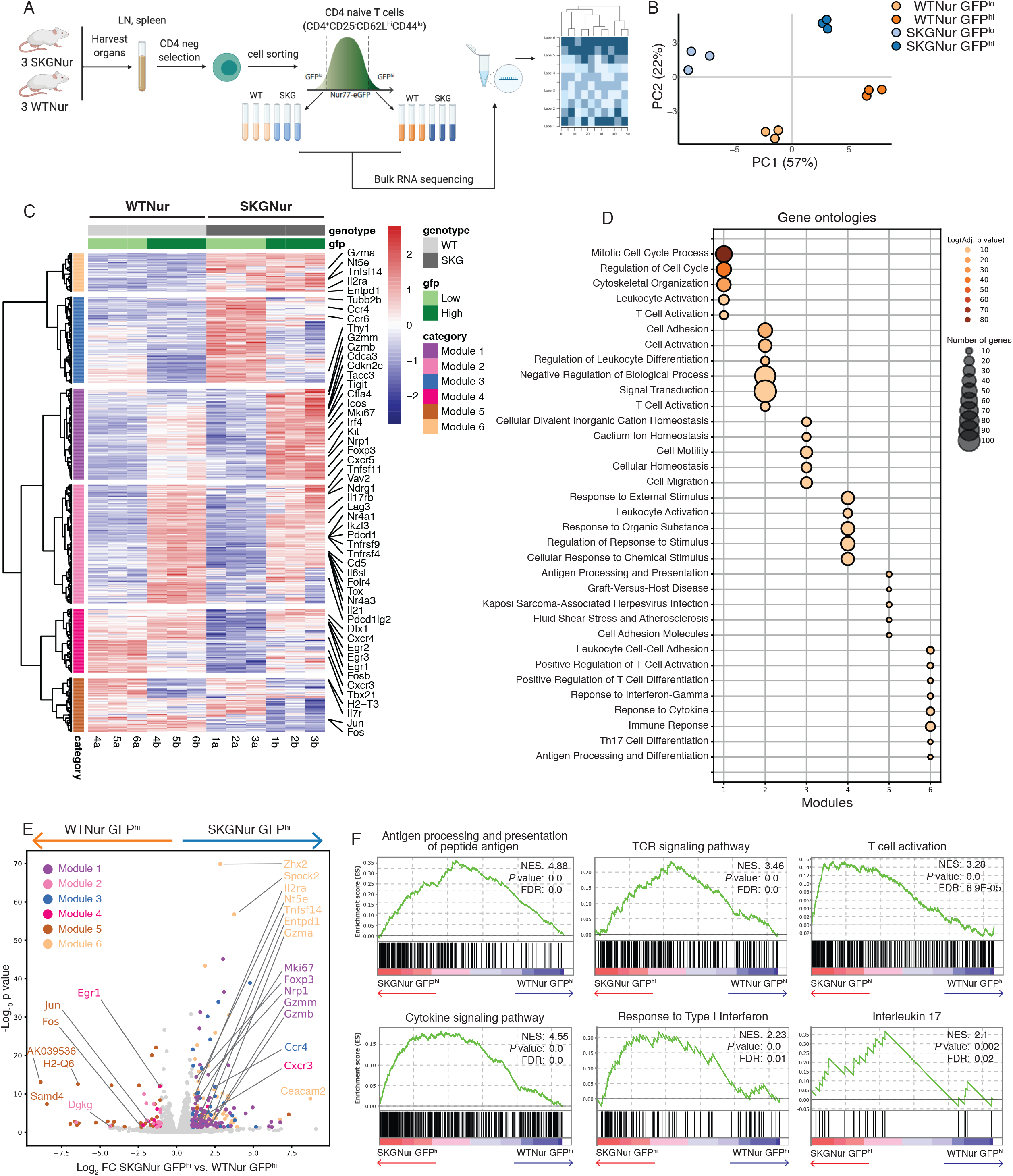
Pre-arthritic naïve SKG T cells demonstrate enhanced T cell activation. (**A**) Experimental scheme of bulk RNA-seq analysis. (**B**) Principal component analysis (PCA) based on transcriptomic data from bulk RNA-seq data shows distribution of SKGNur and WTNur GFP^lo^ and GFP^hi^ CD4 naïve T cell subsets as shown in (A) (*n*=3 per subgroup). (**C**) Heatmap showing expression of 991 differentially expressed genes (DEGs, absolute value(log2(fold-change)) > 1, adjusted *P value <* 0.05) from pairwise comparisons for all samples grouped by subgroup. Hierarchical clustering was used to group DEGs into 6 modules (indicated by dendrogram and row annotation color bar on left). (**D**) Dotplot of select pathways from gene ontology analysis for each gene module from (C) with dot color indicating adjusted *P* value and dot size proportional to number of genes in overlap between pathway genes and module genes. (**E**) Volcano plot of DEGs for SKGNur GFP^hi^ versus WTNur GFP^hi^. DEGs are colored by module membership from gene modules in (C). (**F**) Enrichment plots of TCR signaling and cytokine pathways from GSEA analysis of all GO:BP pathways for ranked genes from SKGNur GFP^hi^ versus WTNur GFP^hi^ differential expression analysis. FDR, false discovery rate. NES, normalized enrichment score

To test this hypothesis, we studied the TCR repertoire and transcriptome of arthritogenic CD4 T cells in SKG mice by leveraging both bulk and single cell RNA sequencing. We capitalized on the SKGNur model in order to capture arthritogenic cells <u>*before*</u> disease onset (akin to the pre-RA phase of disease (18)). We reasoned this could reveal early events in pathogenesis and identify novel targets to preserve tolerance and prevent disease. Within naïve CD4 SKG T with high arthritogenic potential, we found a cluster of cells with the highest expression of *Nr4a1* that upregulate TCR-dependent gene expression programs and exhibit evidence of chronic antigen-stimulation in the pre-disease state. Though these arthritogenic SKG T cells also express anergy-associated genes, we identified a defect in the extent to which they do so relative to WT CD4 T cells. Furthermore, simultaneous determination of SKGNur^hi^ T cell TCR sequences associated with these arthritogenic, chronically antigen-stimulated CD4 T cells in SKG mice revealed an enrichment of naïve CD4 T cells with specific TCR variable beta (Vβ) chains that are known to recognize endogenous mouse mammary tumor viral (MMTV) superantigen in BALB/c mice. We had previously shown that these Vβs escape negative selection in SKG mice (13). Here we find that T cells bearing these Vβs are strongly associated with an activated TCR signaling program in the periphery. We confirmed enrichment of these *TRBV*s in SKG naïve CD4 GFP^hi^ T cells by TCR Vβ protein expression as well. Moreover, the frequency of these CD4 T cells bearing Vβs that recognize MMTV superantigen are expanded in the arthritic joints of SKG mice and likely contribute to development and/or severity of arthritis. Our results reveal how self-reactive SKG T cells that escaped negative selection harbor an independent defect in peripheral tolerance that, together with chronic antigen stimulation, sets the stage for disease. Moreover, we propose a novel role for endogenous MMTV superantigen in promoting arthritogenic T cell responses.

## Results

### Arthritogenic SKG naïve CD4 T cells have a uniquely activated phenotype

We recently demonstrated that it is possible to identify autoreactive and arthritogenic naïve CD4 T cells prior to disease induction on the basis of Nur77-eGFP reporter expression (**Supplementary Fig. 1A**) (11). To understand the transcriptional program of the most arthritogenic CD4 T cells (the SKGNur GFP^hi^ population) prior to disease onset, we performed bulk RNA-sequencing from SKG and wild-type control (SKGNur and WTNur) mice. We sorted and sequenced the 10% highest (GFP^hi^) and 10% lowest (GFP^lo^) Nur77-eGFP-expressing naïve (CD62L^hi^CD44^lo^CD25^-^) CD4 T cells to compare transcriptomes between these four subgroups (**Fig. 1A**, **Supplementary Fig. 1B, Supplementary Data 1**). Principal component (PC) analysis revealed all four subgroups are transcriptionally distinct with the largest variance explained by PC1 (57%) which separated WT and SKG samples by GFP expression followed by PC2 (22%) which separated samples by genotype (**Fig. 1B**). Hierarchical clustering of the collection of 991 differentially expressed genes (DEGs) from subgroup comparisons identified six gene modules that capture the transcriptional differences between the four subgroups (**Fig. 1C**, **Supplementary Data 2**). Gene ontology analysis (19) revealed functional heterogeneity between, and in some cases within, these modules whose expression patterns revealed the unique transcriptomic signature of the SKGNur GFP^hi^ cells. Both SKGNur subgroups (GFP^hi^ and GFP^lo^) uniquely upregulated genes associated with cytokine signaling, antigen processing, and Th17 differentiation (represented in module 6) (**Fig. 1C-D**). Consistent with this, we recently showed that SKG naïve CD4 T cells are hyperresponsive to IL-6 stimulation and produce IL-17 (11). Despite the impaired *in vitro* TCR signaling capability of the more arthritic SKGNur GFP^hi^ subgroup (11), the SKGNur GFP^hi^ and WTNur GFP^hi^ subgroups both distinctly upregulated TCR signaling responsive genes including *Nr4a1, Nr4a3, CD5, Tnfrsf9, Tnfrsf4, Irf4, Tigit, Tox, Pdcd1, Lag3, Ctla4* (found in modules 1 and 2, **Fig. 1C-D**). To further dissect the differences in WTNur GFP^hi^ and SKGNur GFP^hi^ transcriptomic profiles, we focused on the 260 DEGs between these two subgroups. (**Fig. 1E**). Cell cycle and some T cell activation gene programs (e.g., *Cdca3, Cdk2nc, Mki67*; and *Irf4, Ctla4*, *Ikzf2*, *Slamf6, Birc5, Nrp1*, *Tnfsf14* primarily represented in module 1), were more highly upregulated in the SKGNur GFP^hi^ subgroup; whereas genes associated with signal transduction and the negative regulation of a biologic process (e.g., *Nr4a1, Cd5, Tnfrsf9, Folr4/Izumo1r* seen in module 2) were enriched in the WTNur GFP^hi^ subgroup (**Fig. 1C-E, Supplementary Fig. 1C**). Thus, while both GFP^hi^ groups were associated with increased TCR signaling and activation, our data suggests WTNur GFP^hi^ cells may more efficiently induce negative regulators of TCR signaling (module 2).

In an unbiased orthogonal approach, we performed gene set enrichment analysis (GSEA) (20, 21) on the ranked gene list of DEGs and found enrichment of genes upregulated in the SKGNur GFP^hi^ subgroup compared to WTNur GFP^hi^ for pathways associated with antigen processing, T cell signaling and activation, as well as cytokine signaling, including interferon (IFN) responsiveness and IL-17 production (**Fig. 1F**, **Supplementary Data 3**). The SKGNur GFP^hi^ subgroup have a severe impairment in TCR signaling capacity due to the SKG hypomorphic *Zap70* allele (10, 11, 13), and yet they paradoxically upregulated transcriptional TCR and cytokine signaling programs. This likely reflects increased tonic signaling in the SKGNur GFP^hi^ subset due to chronic antigen encounter in the setting of a more autoreactive repertoire. However, these SKG CD4 T cells appeared to have a defect in the ability to efficiently upregulate the usual negative regulators of TCR signaling (e.g., *Folr4*, also known as *Izumo1r)* compared to their induced expression in WTNur GFP^hi^ cells (11, 22–25).

### Pronounced transcriptional heterogeneity among naïve peripheral CD4 T cells

The long half-life of eGFP (26) compared to the more dynamic turnover of NUR77/*Nr4a1* protein and transcript (27–29) (**Supplementary Fig. 2A-C**), meant cells in the GFP^hi^ subgroups most likely represented a heterogeneous collection of more and less recently stimulated cells. To overcome this limitation and to understand whether the TCR signaling and the effector cytokine modules that we identified were uniformly or heterogeneously activated among the subgroups, we performed paired single-cell RNA and TCR sequencing (scRNA- and scTCR-seq) on GFP^hi^ and GFP^lo^ naïve CD4 T cells from SKGNur and WTNur mice (**Fig. 2A**). Analysis of the scRNA-seq dataset revealed significant heterogeneity within the unperturbed, naïve CD4 T cell compartment (defined by surface markers CD62L^hi^CD44^lo^CD25^-^) (**Fig. 2B-C**) across all the subgroups, consistent with a recent report (30). We identified eight distinct clusters, including a cluster (T.N4*_Nr4a1_*) uniquely representing cells that expressed both *Nr4a1* (log2FC=6.1, *P*<2E-308, Wilcoxon rank-sum) and *eGfp* (log2FC=6.0, *P*<2E-308) (**Fig. 2C, Supplementary Fig. 2D, Supplementary Data 4**). The transcriptomic signatures of these eight clusters recapitulated and further refined our bulk RNA-seq gene signatures (**Fig. 1**, **Supplementary Data 5**). Indeed, genes up-regulated in T.N4*_Nr4a1_* cells were overwhelmingly enriched for T cell activation and TCR signaling response genes *(Nr4a1, Nr4a3, Cd5, Tnfrsf9, Tnfrsf4, Egr1-3, Izumo1r, Ifr4*, **Supplementary Data 5**) and primarily overlapped with module 2 (signal transduction and the negative regulation of a biologic process) and module 4 (response to external stimuli and leukocyte activation) genes identified in our bulk RNA-seq analysis (**Fig. 1C-D, Supplementary Fig. 2G, Supplementary Data 2**).

**Figure 2.**
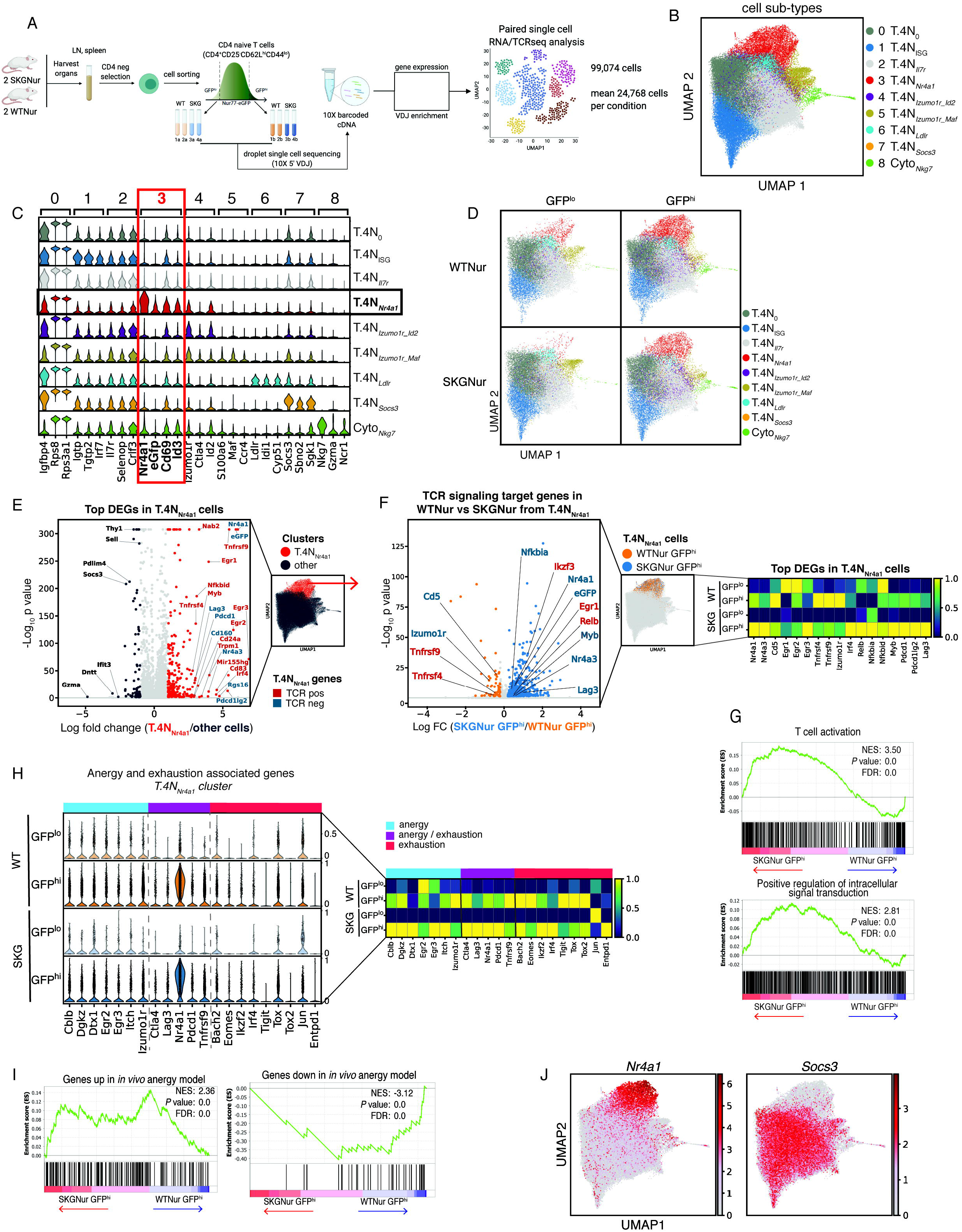
Single-cell RNA sequencing unveils heterogeneity among naïve CD4 T cells with a subset marked by genes associated with TCR signaling. (**A**) Experimental scheme of paired scRNA- and TCR-seq of sorted GFP^hi^ and GFP^lo^ naïve (CD62L^hi^CD44^lo^CD25^−^) CD4 T cells. (**B**) Uniform manifold approximation and projection (UMAP) of 99,074 naïve T cells derived from 8 samples (2 replicates from each WTNur and SKGNur GFP^lo^ and GFP^hi^ subset). Cells are colored according to annotated leiden clusters. (**C**) Stacked violin plot of expression of marker genes for each annotated cluster. Red and black boxes highlight genes uniquely expressed in T.4N*_Nr4a1_* cluster. (**D**) UMAP of cells separated by subgroup (WTNur and SKGNur GFP^lo^ and GFP^hi^ subgroups). (**E**) Volcano plot of cells’ DEGs from T.4N*_Nr4a1_* cluster versus other cells. Dots are colored by significant overexpression (absolute value(log2(fold-change)) > 1, adjusted *P value <* 0.05) in T.4N*_Nr4a1_* cluster (red), other cells (dark grey), or no significant difference (light grey). Labelled genes are colored by role in regulation of TCR signaling either positive (red) or negative (blue). (**F**) Volcano plot of T.4N*_Nr4a1_* cells’ DEGs from SKGNur GFP^hi^ versus WTNur GFP^hi^. Dots are colored as significantly overexpressed (adjusted *P value <* 0.05) in WTNur GFP^hi^ (orange), SKGNur GFP^hi^ (blue), or not significantly different between groups (grey). Labelled genes involved in TCR signaling are colored as indicated in (E). Heatmap shows average expression of the same genes by subgroup normalized by standard scale by column (subtract minimum and divide by maximum). (**G**) Enrichment plots of TCR activation and signaling pathways from GSEA analysis of all GO:BP pathways for ranked genes from differential gene expression analysis of cells in T.4N*_Nr4a1_* cluster from SKGNur GFP^hi^ versus WTNur GFP^hi^. FDR, false discovery rate. NES, normalized enrichment score. (**H**) Stacked violin plot of expression of candidate anergy and exhaustion associated genes in WTNur and SKGNur GFP^lo^ and GFP^hi^ CD4 naïve cells from T.4N*_Nr4a1_* cluster. Heatmap shows average expression of the same genes by subgroup normalized by standard scale by column (subtract minimum and divide by maximum). (**I**) Enrichment plots of gene signature for *in vivo* naïve CD4 anergy model (Ngyugen et al. 2021) for ranked genes from differential gene expression analysis of cells in T.4N*_Nr4a1_* cluster from SKGNur GFP^hi^ versus WTNur GFP^hi^. FDR, false discovery rate. NES, normalized enrichment score. **(J)** UMAP of all cells colored by expression of the indicated genes. Scale is for the log-normalized gene expression.

The GFP^hi^ T cells from both SKGNur and WTNur mice were clearly enriched in the T.N4*_Nr4a1_* cluster compared to GFP^lo^ CD4 T cells in the respective mice by a mean of > 4-fold. GFP^hi^ T cells were also enriched in the T.N4*_Izumo1_Id2_* and, albeit to a lesser extent, the Cyto*_Nkg7_* clusters (**Fig. 2D; Supplementary Fig. 2D-E**) but were present across all eight clusters. This suggested to us that only a fraction of the GFP^hi^ naïve CD4 T cells had more recent antigen encounter, whereas most of the other GFP^hi^ T cells had downregulated their *Nr4a1* and *eGfp* transcripts and segregated into heterogeneous phenotypes. We next closely examined the T.N4*_Nr4a1_* cluster to determine which transcriptional changes in this cluster primed SKGNur GFP^hi^ naïve CD4 T cells to differentiate into arthritogenic T cells.

### A subset of GFP^hi^ naïve T cells is transcriptionally defined by an enhanced TCR signaling program

Given the specificity of NR4A1 (NUR77) as a reporter of recent TCR signaling (31), the high expression of *Nr4a1* in the T.N4*_Nr4a1_* cluster signifies that these cells were actively, or at least recently, receiving TCR signaling input, most likely due to endogenous antigen(s) encounter (23, 24). The T.N4*_Nr4a1_* cluster highly expressed multiple TCR response genes (*Nr4a1, Egr1-3, Irf4, Irf8, Tnfrsf9, Nfkbid, Tnfrsf4, Bcl2a1b, Myb, Ikzf2, Nfkb2, Cd69*), as well as negative checkpoint regulators (*Pdcd1Ig2, Pdcd1, Lag3)* (**Fig. 2E**, **Supplementary Data 4**) that are known to be induced by T cell activation to fine-tune and restrain T cell responses, enforce peripheral tolerance, and limit immunopathology (32–35). This TCR signaling signature was remarkably further upregulated in the signaling impaired GFP^hi^ T cells from SKGNur mice within the T.N4*_Nr4a1_* cluster (**Fig. 2F-G**, **Supplementary Data 3 and 5**), thereby mirroring the enrichment for genes involved in TCR signaling and T cell activation in our bulk RNA-seq dataset (**Fig. 1F, Fig. 2G**). Our scRNA-seq findings suggest active antigen encounter and adaptive gene regulation primarily occurred in the T.N4*_Nr4a1_* cells and to a greater extent in the SKGNur GFP^hi^ cells within this cluster. We next investigated additional T cell transcriptomic signatures that could further illuminate how SKGNur GFP^hi^ cells in the T.N4*_Nr4a1_* cluster are poised to escape tolerance.

### Endogenous antigen-activated T cells upregulate a tolerogenic transcriptional program

T cell-intrinsic mechanisms that operate during thymic development (negative selection of self-reactive cells) and in peripheral T cells (functional unresponsiveness or ‘anergy’) are essential to maintain tolerance to self. Therefore, we sought to examine the expression of candidate genes known to be associated with these two tolerance programs (36–41) in SKGNur and WTNur GFP^hi^ cells compared to their GFP^lo^ counterparts within the T.N4*_Nr4a1_* cluster, which had the highest expression of *Nr4a1* and *Nr4a3* (log2FC=4.0, adjusted *P*<1E-42). We found that in the T.N4*_Nr4a1_* cluster, and in the overall dataset, GFP^hi^ naïve CD4 T cells upregulated anergy and exhaustion-associated gene modules in both the WTNur and SKGNur subgroups (**Fig. 2H, Supplementary Fig. 2H**). Additionally, the arthritogenic SKGNur GFP^hi^ naïve CD4 T cells within the T.N4*_Nr4a1_* cluster upregulated genes associated with a gene signature enriched in an *in vivo* anergy model compared to WTNur GFP^hi^ naïve CD4 T cells (25) (**Fig. 2I**). These results coincide with our published work defining a subset of Nur77-eGFP high CD4 T cells with a naïve surface phenotype juxtaposed with evidence of self-reactivity and upregulation of anergic markers (11, 25, 44). This likely reflects the triggering of a negative regulatory program in naïve CD4 T cells in response to tonic TCR signaling since NR4A family members have been shown to play negative regulatory roles in peripheral T cells (28, 41–43), and to be associated with transcriptional signatures of anergy and exhaustion including expression of *PDCD1* (PD-1), *TIGIT*, and other inhibitory regulators (41).

However, within the SKGNur GFP^hi^ cells, upregulation of these tolerogenic gene expression patterns appeared insufficient to induce anergy. Indeed, although genes upregulated in SKGNur GFP^hi^ T cells are associated with an anergy gene signature (**Fig. 2H-I**), Folate receptor 4 (FR4) gene expression (*Izumo1r*) – a marker of anergic cells – was significantly lower in SKGNur GFP^hi^ compared to WTNur GFP^hi^ T cells (**Fig. 2H**, **Supplementary Fig. 2H**). This is consistent with our previous report of SKGNur mice in which we described a reduced frequency of anergic peripheral T cells (11). Therefore, it appears that the anergy program is likely defective in SKG CD4 T cells, perhaps due in part to their severely impaired proximal TCR signaling capacity. Therefore, in addition to a known loss in central tolerance, SKG mice have an independent defect in peripheral tolerance with unchecked activation of arthritogenic clones that cause autoimmune, erosive arthritis.

### SKG’s hyperresponsiveness to IL-6 is pre-programmed transcriptionally

We previously found that SKGNur GFP^hi^ T cells were more responsive to IL-6 and more readily produced IL-17 in the most autoreactive T cells, in part due to lower levels of SOCS3 (suppressor of cytokine signaling 3) – a critical negative regulator of IL-6 (11) (**Supplementary Fig. 1A**). Here, we found that genes associated with IL-6 signaling machinery and the Th17 pathway were uniquely enriched in SKGNur GFP^hi^ T cells (45) in the T.N4*_Nr4a1_* cluster (**fig. S2I**), consistent with our bulk RNA-Seq results (**Fig. 1F**, **Supplementary Data 3**).

SOCS3 is downregulated in naïve CD4 T cells in response to antigen (46) and in patients with RA (11, 47). Its expression has a strong inverse correlation with murine arthritis severity (48–50). Therefore, we examined the expression of SOCS family members in our single cell dataset. Of these family members, *Socs3* was specifically downregulated in SKGNur GFP^hi^ cells within the T.N4*_Nr4a1_* cluster (**Supplementary Fig. 2J**). Moreover, we found a striking inverse correlation between individual T cells that expressed *Nr4a1* and *Socs3* (**Fig. 2J**), corresponding to a published report (46). The stark inverse correlation between *Nr4a1* and *Socs3* expression in SKGNur GFP^hi^ T cells within the T.N4*_Nr4a1_* cluster provides orthogonal validation of our previous results. It highlights the interdependence between signaling via the TCR and heightened sensitivity to cytokines such as IL-6 (11), providing yet another link between T cell self-reactivity and pathogenicity.

### T.N4_Nr4a1_ cells segregate into two distinct TCR signaling modules

To complement our investigation of these curated pathways that could contribute to pathogenicity of SKGNur GFP^hi^ T cells in the T.N4*_Nr4a1_* cluster, we turned to unbiased methods for discovery of transcriptomic signatures within the T.N4*_Nr4a1_* cluster to elucidate additional unknown mechanisms. We identified three distinct modules of highly variable genes (HVGs) that positively correlated with *Nr4a1* (**Fig. 3A**). To our surprise, we found that members of two of these modules, *Egr* family members (immediate early gene transcription factors) and *Tnfrsf9* (4-1BB – the TCR inducible co-stimulatory receptor), identified distinct subsets of cells within the T.N4*_Nr4a1_* cluster (**Fig. 3B** and **Supplementary Fig. 3A**). Indeed, within just the T.N4*_Nr4a1_* cells, *Egr’s* and *Tnfrsf9* continued to segregate into separate gene modules of HVGs positively correlated with *Nr4a1* (**Supplementary Fig. 3B**). The *Egr* module contained additional immediate early genes or markers of early T cell activation (e.g., *Egr1, Egr2, Cd69, Ier2, Egr3, Nfkbid, Junb, Fos, Myc, Cd40lg*), whereas the *Tnfrsf9* module contained markers of chronic or prolonged TCR signaling input (e.g., *Nr4a1*, *Pou2f2, Myb, Tnfrsf4, Lag3*) (**Fig. 3C** and **Supplementary Fig. 3C**, **Supplementary Data 6**).

**Figure 3.**
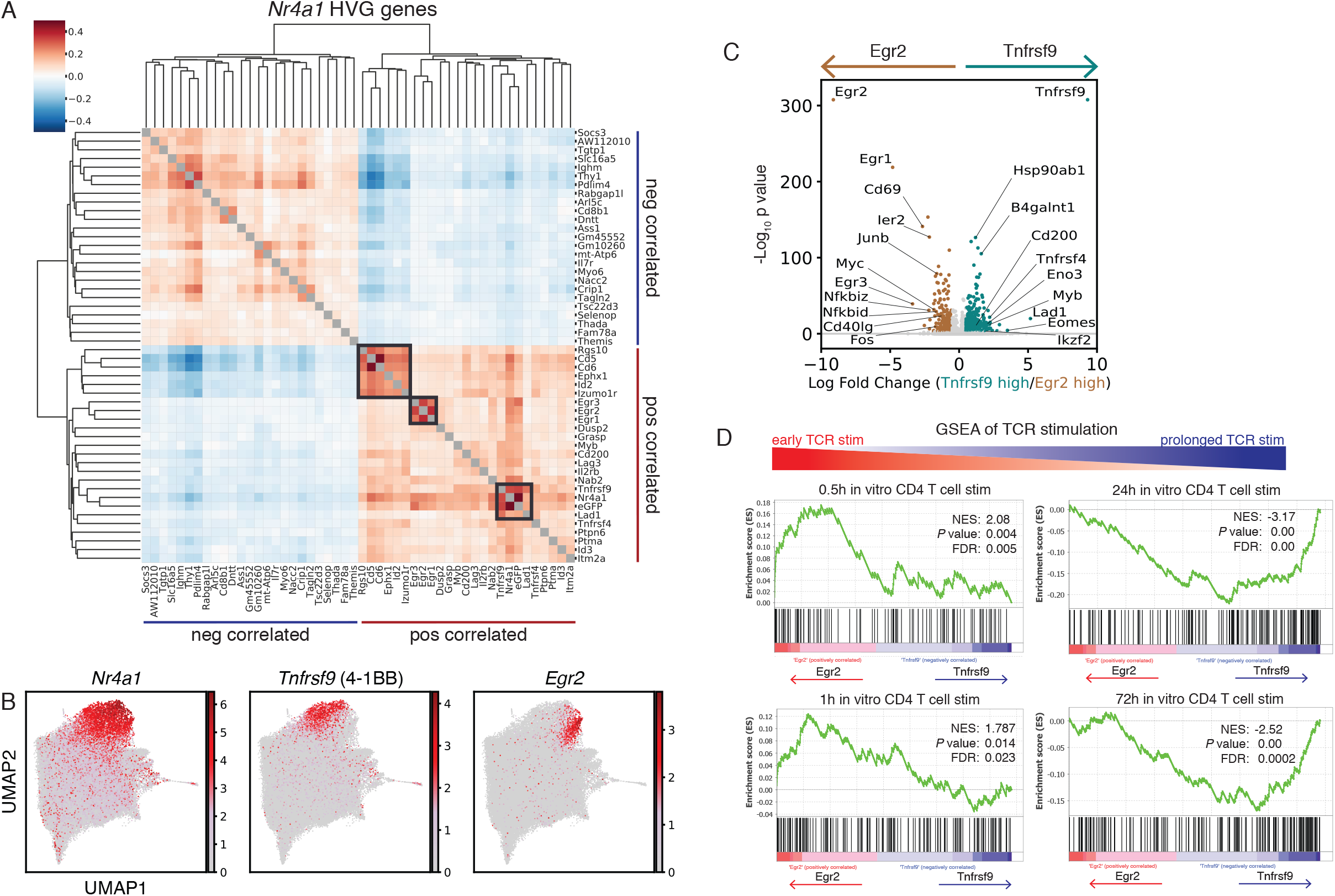
T.N4_*Nr4a1*_ cells segregate into two distinct TCR signaling modules that segregate acute from chronic antigen-activated T cells. (**A**) Correlation matrix shows hierarchical clustering of Spearman’s correlation of top 25 highly variable genes (HVG) that positively and negatively correlate with *Nr4a1* expression across all cells. Diagonal grey boxes represent correlation of 1. Dark grey boxes mark distinct gene modules from genes that positively correlate with *Nr4a1* expression. (**B**) UMAP plots show the expression levels of indicated marker genes. Scale represents the log-transformed normalized counts of genes. (**C**) Volcano plot shows DEGs for cells in T.4N_Nr4a1_ cluster that expressed (log normalized expression > 1) *Egr2* or *Tnfrsf9* with dots colored by significant overexpression (absolute value(log2(fold-change)) > 0.05, adjusted *P value <* 0.05) in *Egr2* (tan) or *Tnfrsf9* (teal) expressing cells. (**D**) Enrichment plots of pathways of time course *in vitro* activation of CD4+ T cells with αCD3 + CD28 from GSEA analysis of pathways from study GSE17974 for ranked genes from differential gene expression analysis of cells in T.4N*_Nr4a1_* cluster that express *Egr2* versus *Tnfrsf9*. FDR, false discovery rate. NES, normalized enrichment score.

Using GSEA we found that the genes overexpressed in *Egr2* high expressing cells were associated with gene pathways induced early after TCR stimulation (0.5h and 1h timepoints), whereas *Tnfrsf9* high expressing cells overexpressed genes associated with gene pathways upregulated after prolonged TCR stimulation (24h and 72h, **Fig. 3D**, **Supplementary Data 3**). Our findings suggest that T.N4*_Nr4a1_* cells segregate into subclusters driven by acute versus chronic TCR signaling signatures.

To examine other co-variates which could lead to segregation of acute versus chronic antigen-activated T cells, we performed cell-cycle analysis on our dataset. Though cell-cycle appeared to contribute somewhat to the heterogeneity among the T.N4*_Nr4a1_* cluster, as *Tnfrsf9* positive expressing cells were more likely to be associated with S-phase cell-cycle genes (**Supplementary Fig. 3D**), it did not fully account for the clear division between the expression of the *Egr* family members and *Tnfrsf9.*

### Cell states and trajectories of T.4N_Nr4a1_ cells have a distinct distribution in SKGNur GFP^hi^ subset

We next investigated if these acute versus prolonged TCR signaling states were related and may contribute to the arthritogenic potential of the SKGNur GFP^hi^ subset. Specifically, we asked if the early vs prolonged TCR signaling signatures associated with *Egr2* and *Tnfrsf9* markers, respectively, were indicative of a continuum of cell states. We used RNA velocity analysis (51) to create a latent time axis for cells from the T.N4*_Nr4a1_* cluster and asked if this latent time axis related to the acute versus chronic TCR signaling states delineated by *Egr2* and *Tnfrsf9*. Qualitatively, cells assigned an earlier latent time appeared to overlap with *Egr2* high cells in the UMAP, whereas cells assigned a later latent time overlap with *Tnfrsf9* expression (**Fig. 3B, 4A**). Additionally, the expression of *Egr2* and associated genes peaked in cells earlier in the latent time axis, while the expression of *Tnfrsf9* and associated genes peaked in cells later along the latent time axis (**Fig. 4B-C**).

**Figure 4.**
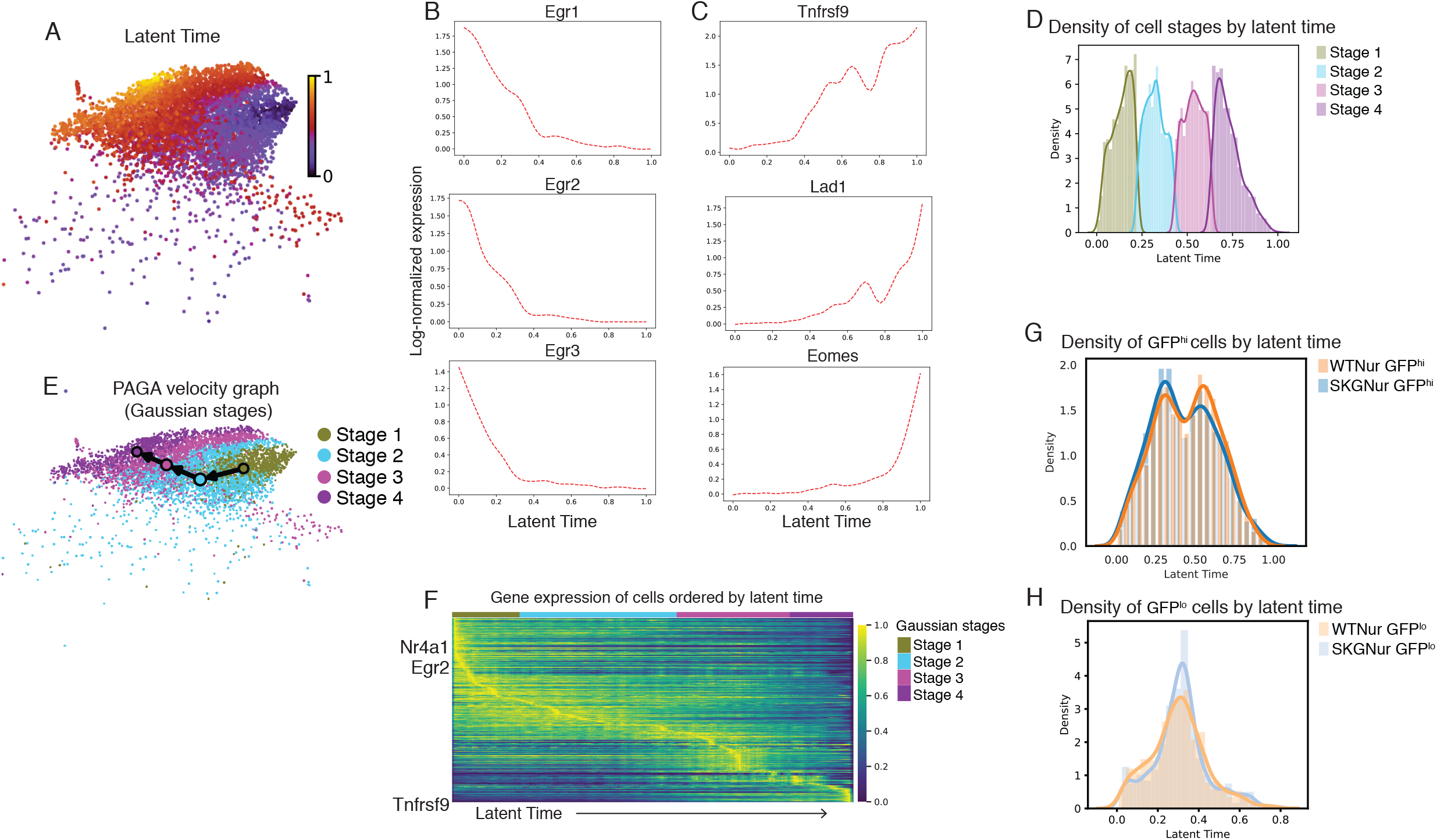
Trajectory analysis of T.4N_Nr4a1_ cells orthogonally uncovers acute versus chronic antigen-activated T cell states with a distinct distribution in the SKGNur GFP^hi^ subset. (**A**) UMAP of cells from T.4N_Nr4a1_ cluster colored by latent time assigned by the scvelo dynamical modeling algorithm. (**B**) Smoothed gene expression from cells in T.4N_Nr4a1_ cluster of select genes with highest expression earlier or later **(C)** along latent time axis. (**D**) Probability densities of latent time distribution of cells from T.4N_Nr4a1_ cluster assigned to 4 distinct clusters (labelled “Stage 1” – “Stage 4”) by a Gaussian mixture model. (**E**) Predicted transitions from PAGA algorithm between cells from stages indicated in (D). (**F**) Single cell heatmap of standard scale normalized expression of genes ordered top to bottom by peak expression at earlier to later latent time. Chosen genes are the top 300 genes with the highest confidence used in modeling of latent time. Column annotation bar indicates stage assignment of the cell in each column. (**G-H**) Probability densities of latent time distribution for GFP^hi^ (**G**) and GFP^lo^ (**H**) cells from WTNur and SKGNur mice.

We next used an unbiased deconvolution method to separate the bimodal distribution of all cells across the latent time axis into 4 underlying distributions or cell states labelled “Stage 1” to “Stage 4” from earlier to later latent time (**Fig. 4D**, **Supplementary Fig. 3E-F**). The UMAP overlay of the dynamical model velocity vector field qualitatively suggests sequential transitions between the stages (**Supplementary Fig. 3G**). These transitions were further supported by trajectory inference analysis (52) which predicted a trajectory from Stage 1 to Stage 4 (**Fig. 4E**). The expression of *Egr2* and *Nr4a1* peaked within cells from Stage 1 while the expression of *Tnfrsf9* peaked within cells from Stage 4 (**Fig. 4F, Supplementary Data 7**), and the genes overexpressed in Stage 1 versus Stage 4 cells were enriched for the same pathways induced after early or after prolonged TCR stimulation as had been enriched in the *Egr2* high and *Tnfrsf9* high cells, respectively (**Supplementary Fig. 3H**). Thus, our RNA velocity analysis independently uncovered the same cell states defined by the *Egr2* and *Tnfrsf9* marker genes and suggests that these cell states are the endpoints of a trajectory of cell states from early to prolonged TCR stimulation.

We then asked if there were any differences between the distribution of cells from the SKG and WT subgroups among these cell states. Cells from SKGNur GFP^hi^ and WTNur GFP^hi^ groups had significantly different bimodal distributions across latent time with a higher density at earlier time for the SKGNur GFP^hi^ cells and a higher density at later time for the WTNur GFP^hi^ cells, which was not observed in the GFP^lo^ subsets (**Fig. 4G-H**) (p = 0.002 and p = 0.11, respectively, from the Kolmogorov-Smirnov test). The cells from SKGNur GFP^hi^ had an increased odds of being in Stage 1 versus Stage 4 than the cells from WTNur GFP^hi^ which agreed with our latent time density for each of those cell groupings (OR = 1.25, p = 0.02). We hypothesized that the arrest of SKGNur GFP^hi^ cells from the T.N4*_Nr4a1_* cluster in Stage 1, which is associated with early TCR stimulation, was not due to these cells truly experiencing primarily acute TCR stimulation but instead that prolonged TCR stimulation within these SKGNur GFP^hi^ cells did not robustly induce the Stage 4 prolonged TCR transcriptomic signature as seen for the WTNur GFP^hi^ cells. These results correspond to our bulk RNASeq findings (**Fig. 1C-D**) and support our hypothesis that SKG CD4 T cells have a defect in peripheral tolerance induction resulting in a reduced frequency of anergic cells (11).

### Arthritogenic naïve CD4 T cells in SKG mice demonstrate a biased TCR variable beta gene repertoire

We have previously shown that the NUR77 reporter of TCR signaling can enrich for arthritogenic naïve (CD62L^hi^CD44^lo^) T cells characterized in part by their increased autoreactivity and ability to proliferate in response to an undefined endogenous antigen(s) (11). Thus, we asked how the SKGNur GFP^hi^ TCR repertoire might be contributing to the activation of these T cells in the periphery. We examined the TCR repertoire in the subsets using scTCR-seq (**Fig. 2A**) and detected paired TCR α (TRA) and TCR β (TRB) genes in 86% of cells (**Supplementary Fig. 4A**). Using Gini coefficient analysis, we did not find an oligoclonal expansion in any of the clusters of naïve T cells sorted from any of the samples, including all the SKGNur samples (**Supplementary Data 8**).

Instead, we found that SKGNur GFP^hi^ T cells are uniquely associated with biased TCR variable β gene (*TRBV*) usage, but not TCR variable α gene (*TRAV*) usage (**Fig. 5A-C**, **Supplementary Fig. 4B**). In SKGNur GFP^hi^ CD4 T cells compared to the paired SKGNur GFP^lo^ samples, we found significantly higher (FDR < 0.1) usage of *TRBV26* (TCR variable beta (Vβ) 3), *TRBV12-1* (Vβ5), *TRBV15* (Vβ12), *TRBV16* (Vβ11), *TRBV3*, and *TRBV29* (Vβ7) and each of these *TRBV* genes also had a higher mean frequency in SKGNur GFP^hi^ cells compared to WTNur GFP^hi^ cells (**Fig. 5A,C-D**, **Supplementary Data 9**).

**Figure 5.**
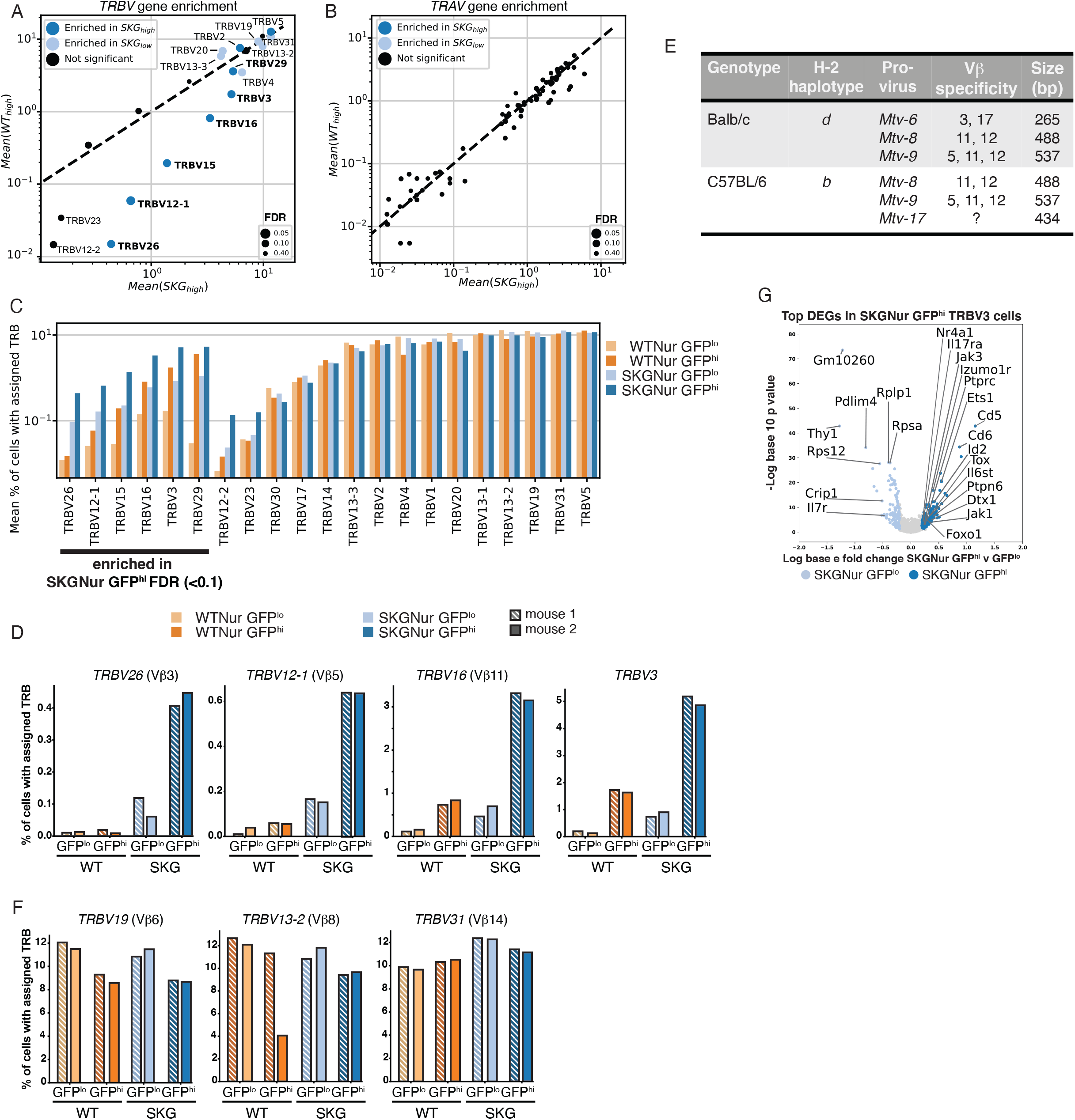
SKG CD4 T cells harbor a biased TCR variable beta gene repertoire. (**A-B**) Scatterplot of mean frequency of cells expressing each TRBV (**A**) or TRAV (**B**) gene for the SKGNur GFP^hi^ samples versus the WTNur GFP^hi^ samples. Dots for each TRBV and TRAV genes are sized according to the FDR from a one-sided paired t-test comparing frequency in SKGNur GFP^hi^ versus SKGNur GFP^lo^. Dots are colored as either significantly enriched (FDR < 0.1) in SKGNur GFP^hi^ (dark blue), significantly enriched in SKGNur GFP^lo^ (light blue), or not significantly enriched in either subgroup (black). Dots for significant TRBV genes are labelled with the TRBV gene name. Labels for TRBV genes that were significantly enriched in SKGNur GFP^hi^ and were also more highly expressed in SKGNur GFP^hi^ samples versus WTNur GFP^hi^ samples are bolded. (**C**) Bar plot of mean value of cells expressing each TRBV gene as a percentage of all cells in each sample with an assigned TRBV. Bars are colored according to subgroup and ordered with the TRBV genes enriched in SKGNur GFP^hi^ (A) followed by the other TRBV genes ordered by increasing overall frequency. (**D**) Bar plots of frequency of cells expressing indicated TRBV genes significantly enriched in SKGNur GFP^hi^ for two replicate mice in each subgroup. (**E**) Table depicting H-2 haplotype, expected Mtv pro-virus, its Vβ specificity and base pair (bp) size on gel for Balb/c and C57BL/6 mice. (**F**) Bar plots of frequency of cells expressing indicated TRBV control genes not uniquely enriched in SKGNur GFP^hi^ cells for two replicate mice in each subgroup. (**G**) Volcano plot of DEGs of cells with assigned indicated TRBV from the SKGNur GFP^hi^ or SKGNur GFP^lo^ subgroups. Dots are colored by significant overexpression (absolute value(natural log(fold-change)) > 0.2, adjusted *P value <* 0.05) in SKGNur GFP^hi^ (dark blue), SKGNur GFP^lo^ (light blue), or no significant difference (light grey).

Polyclonal Vβ expansion occurs in the presence of superantigen in both humans and mice (53, 54). Interestingly, the *TRBV* genes enriched in the SKGNur GFP^hi^ T cell population mark Vβ’s that are known to recognize endogenous viral superantigens from mouse mammary tumor virus (MMTV) (**Fig. 5E**) (55, 56). To further address whether the SKG TRBV repertoire might be shaped by MMTV superantigen, we examined whether our SKG line carry MMTVs previously reported to be present in BALB/c mice (55, 56) and found that indeed *Mtv-6*, *Mtv-8*, *Mtv-9* were present in SKG mice, unlike the *Mtv-17* strain present in C57BL/6 mice (**Supplementary Fig. 4C**). In contrast to the *TRBV* genes uniquely enriched in SKGNur GFP^hi^ cells, *TRBV* genes for Vβ’s not known to respond to these particular MMTV’s (e.g., *TRBV19*/Vβ6, *TRBV13-2*/Vβ8, *TRBV31*/Vβ14) are not enriched in SKGNur GFP^hi^ T cells (**Fig. 5C,F**). These results reinforce the notion that negative selection is defective in SKG mice, particularly in the SKGNur GFP^hi^ T cells, congruent with our previous work (13) and suggests that the biased TRBV repertoire in SKGNur GFP^hi^ cells is indeed likely shaped by encounter with endogenous MMTV superantigen.

Since Nur77-eGFP expression is dynamically expressed in T cells, the observed *TRBV* restriction suggests that the SKG *TRBV* repertoire in GFP^hi^ T cells is driven by involvement of viral superantigen(s) not only in the thymus, but also in the periphery. It is likely that encounter with endogenous viral superantigens drives their *Nr4a1* expression and increased frequency. Indeed within the T.N4*_Nr4a1_* cluster, representing the most recently stimulated cells, SKGNur GFP^hi^ cells also had enrichment of several Vβ’s known to respond to these specific MMTV’s (e.g. *TRBV16* (Vβ11) **Supplementary Fig. 5A-D**). This suggests that there is an intersection between cells occupying the transcriptional states in this cluster which are enriched in TCR signaling target genes and cells with Vβ’s that recognize MMTV superantigen (**Supplementary Fig. 5A-D**). Consistent with this, the subset of cells from the entire SKGNur GFP^hi^ population that express the enriched *TRBV3* and likely recognize MMTV superantigen upregulated a unique profile of TCR signaling target genes more than their GFP^lo^ counterparts expressing the same *TRBV* (**Fig. 5G**). This is not unique to *TRBV3* (**Supplementary Fig. 4D, Supplementary Data 10**).

Taken together, these results strongly suggest that the biased *TRBV* repertoire in SKGNur GFP^hi^ cells is shaped by impaired negative selection and peripheral encounter with endogenous MMTV superantigen, the latter of which is contributing to their more activated state. Indeed, the superantigen recognition in the periphery in SKGNur GFP^hi^ cells within the enriched *TRBV* repertoire could be influencing their unique transcriptional states especially those found within the recently TCR stimulated cells in the T.N4*_Nr4a1_* cluster.

### Arthritogenic T cells are enriched for Vβ’s driven by endogenous superantigen(s)

To validate our scTCR-seq results, we examined the distribution of TCR Vβ protein levels in SKGNur and WTNur peripheral CD4 T cells prior to arthritis initiation using antibody staining against a subset of candidate Vβ’s with commercially available antibodies (gating strategy **Supplementary Fig. 6A**). We found that the Vβ protein abundances paralleled the transcript abundances observed in our scTCR-seq dataset. T cells expressing Vβ3, Vβ5, Vβ11 (corresponding to *TRBV26*, -*12*, -*16* respectively) were significantly enriched in SKGNur GFP^hi^ peripheral naïve CD4 T cells from lymph node (LN) and spleen (**Fig. 6A-B** and data not shown), whereas Vβ’s not known to respond to BALB/c specific MMTVs (e.g., Vβ6, Vβ8, Vβ14 corresponding to *TRBV19*, -*13*, -*31* respectively) were not enriched in SKGNur GFP^hi^ cells (**Supplementary Fig. 6B-C**, **Fig. 5A,C**).

**Figure 6.**
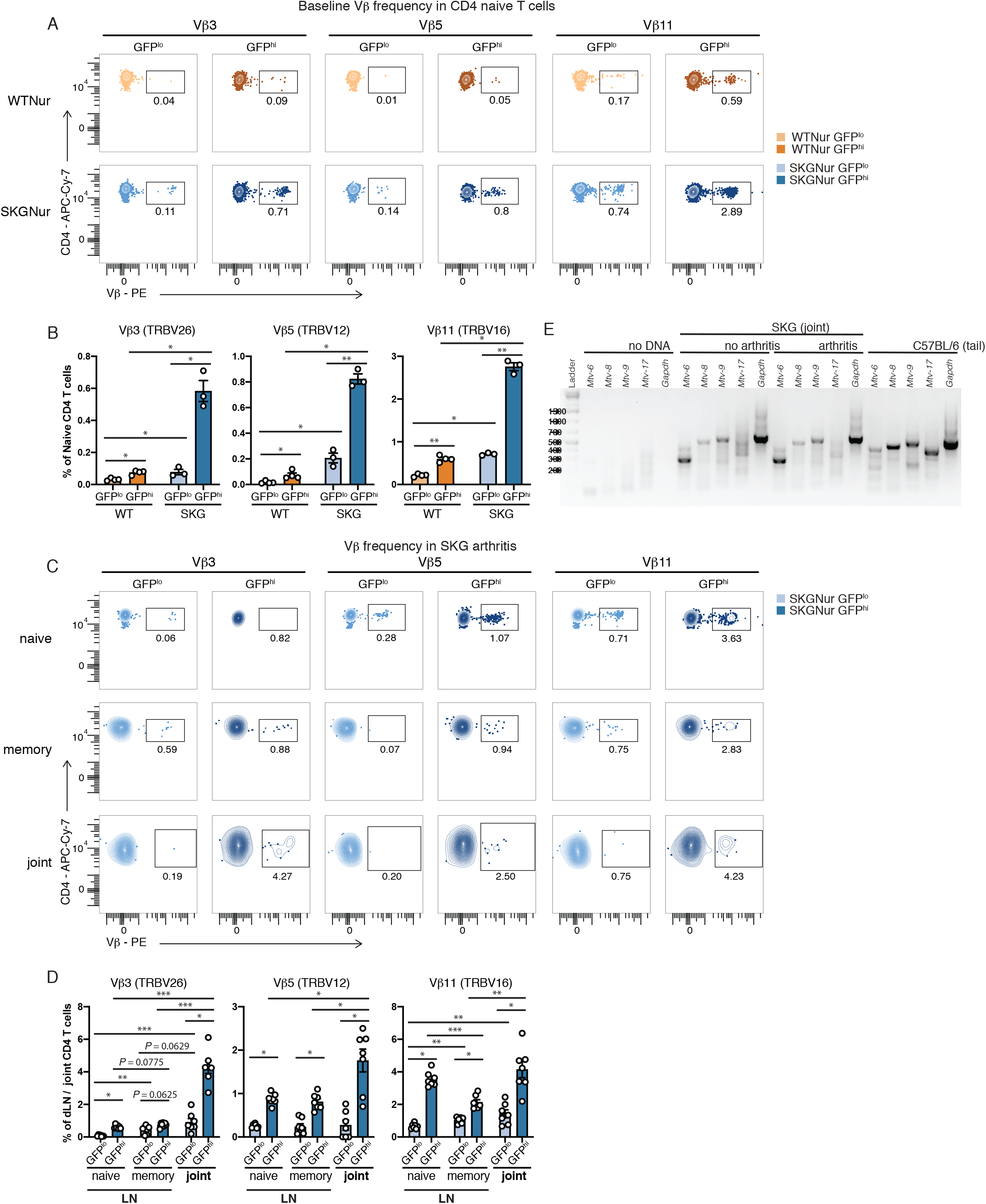
Arthritogenic CD4 T cells are enriched for TCR Vβs likely driven by endogenous superantigen(s). (**A-B**) Representative FACS plots (**A**) of naïve peripheral CD4 T cells with indicated TCR V protein usage determined by flow cytometry in GFP^lo^ and GFP^hi^ T cells from LN of WTNur and SKGNur mice prior to arthritis induction and quantified in (**B**) where bar graphs depict mean frequency (± SEM), n = 3-4 mice per group, experiment repeated at least 3 times. (**C-D**) Representative FACS plots (**C**) of peripheral naïve or memory, or joint CD4 T cells with indicated TCR V protein usage determined by flow cytometry in GFP^lo^ (light blue) and GFP^hi^ (dark blue) T cells from LN or joints of SKGNur mice 2.5 weeks after arthritis induction with zymosan (as seen in Supplemental Fig. 6D) and quantified in (**D**) where bar graphs depict mean frequency (± SEM), n = 7 mice per group pooled from 2 experiments. Significance indicated by asterisk for *P* value (exact permutation test) or FDR (paired t-test) < 0.05 (*), < 0.1 (**), or < 0.001 (***) in (B) and (D). (**E**) DNA was used from SKG joints ± arthritis in PCR reactions containing primers specific for the indicated Mtv pro-viruses. Lanes 2-6 show PCR mixtures lacking template DNA. (D, K) C57BL/6 tail DNA was used as a positive control for *Mtv-8*, −9, −17 and a negative control for *Mtv-*6. Molecular size markers are shown in lane 1. Each gel is representative of at least 3-4 biological replicates per condition and genotype.

We then asked whether there was a further enrichment of the BALB/c MMTV specific Vβ’s in SKG mice in the setting of arthritis. After moderate to severe inflammatory arthritis was established in SKG mice (**Supplementary Fig. 6D**), peripheral CD4 T cells were harvested from joint draining LN (dLN) and arthritic joints and then examined for Vβ usage. We found a significantly higher frequency of Vβ3, Vβ5, and Vβ11 in SKGNur GFP^hi^ T cells compared to T cells from the GFP^lo^ subgroup in both naïve and memory peripheral CD4 T cells, with an even further increase in frequency among SKG GFP^hi^ T cells infiltrating the joint compared to those from the dLN (**Fig. 6C-D**). This further enrichment suggests T cells with these particular Vβ’s encounter antigen in the arthritic joints of SKG mice and is congruent with our previous results that SKG joint infiltrating CD4 T cells upregulate NUR77 in response to antigen encounter (11). This enrichment in the joint was not observed in T cells expressing V 6, V 8, and Vβ14 (**Supplementary Fig. 6E-F**).

These provocative results indicate that Vβ enrichment in the SKGNur GFP^hi^ CD4 T cell subset may be driven by superantigen in the periphery and even the joints. We found that BALB/c specific *Mtv*’s are expressed in the joints of non-arthritic and arthritic SKG mice (**Fig. 6E**). Therefore, presence of endogenous viral superantigen(s) in the joints may lead to engagement of T cells expressing these unique Vβ’s that are transcriptionally poised to enter into an immune response as a result of signaling via their antigen-receptor (marked by *Nr4a1*/NUR77 transcript and protein upregulation) and thus lead to the enrichment of these particular Vβ’s (Vβ3, Vβ5, and Vβ11) in the arthritic joint.

## Discussion

In this study, we leveraged the SKGNur mice and both bulk RNA and single-cell sequencing to profile the gene expression program and TCR repertoire of self-reactive pathogenic T cells before arthritis onset. Using single-cell sequencing, we unmasked remarkable transcriptional heterogeneity in the phenotypically naïve CD4 T cell compartment in both SKG and WT mice. We identified a subset of these cells (T.4N*_Nr4a1_*) that upregulate TCR signaling and TCR activation gene programs in response to recent endogenous antigen encounter (marked by upregulation of *Nr4a1, Cd69, Cd5, Egr1-3, Tnfrsf9, Nfkbid*). We further identify two subsets of naïve CD4 T cells within the T.4N*_Nr4a1_* cluster that were either acutely or chronically antigen-stimulated and enriched for immediate-early gene expression and gene expression associated with prolonged TCR signaling respectively (**Fig. 3B-D, Supplementary Fig. 3H**). These phenotypically naïve CD4 T cells have also upregulated gene modules associated with TCR negative regulators (e.g., *Pdcd1Ig2, Pdcd1, Lag3)* and markers of anergy / exhaustion (*Cblb*, *Dtx1, Egr3, Izumo1r, Ctla4, Lag3, Nr4a1, Tnfrsf9, Tigit, Tox*). Remarkably, many components of these TCR-dependent gene programs are further upregulated in naïve SKGNur GFP^hi^ CD4 T cells compared to WT T cells (**Fig. 1F, 2F-I**) despite their hypomorphic *Zap70* allele, most likely reflective of their more autoreactive repertoire and chronic endogenous antigen encounter.

Despite this chronic exposure to endogenous antigen, naïve SKGNur CD4 GFP^hi^ T cells do not successfully transition to a chronically stimulated transcriptional state, nor do they efficiently transition along the trajectory of cell states associated with early to prolonged TCR stimulation within the T.4N*_Nr4a1_* cluster. These findings may in part explain why SKGNur GFP^hi^ CD4 T cells do not efficiently induce the usual negative regulators of TCR signaling as identified in our bulk RNASeq dataset (**Fig. 1C-D**), unmasking a defect in peripheral tolerance.

Previous examinations of the TCR Vβ repertoire in SKG mice had not identified oligoclonal expansion of a particular Vβ (22). However, our transcriptomic dataset directly probed the TCR-sequences of the more arthritogenic naïve SKGNur GFP^hi^ T cells and revealed repertoire differences at the single cell level. Here we describe a previously unknown TCR Vβ bias in the naïve SKG repertoire. Our results provide a direct link demonstrating how altered thymic selection in SKG mice can result in a biased peripheral *TRBV*/Vβ repertoire within the SKGNur GFP^hi^ subset (**Fig. 5A**,**C,D**, **6A-B**). The Vβ’s enriched in the SKGNur GFP^hi^ subset within our dataset are well known to recognize endogenous mouse viral superantigens (from MMTV, **Fig. 5E**) (55, 57, 58) and suggest that their repertoire is further influenced in the periphery by these superantigen(s). This may have led to a greater degree of basal activity and adaptation of the cells bearing TRBVs/Vβ’s that recognize MMTV superantigen(s) in these mice, thus explaining the higher frequencies of T cells bearing these *TRBVs*/Vβ’s in the SKGNur GFP^hi^ subset. Moreover, the further enrichment of these Vβ’s in the SKG arthritic joint (**Fig. 6C-D**) suggests that the expansion of these Vβ’s may be driven by intra-articular superantigen encounter. These findings may also be relevant in human RA (59–61), and other autoimmune arthritides (62–64), where T cells bearing particular Vβ’s have been reported to be expanded and retained in the synovial microenvironment.

We propose a speculative model drawn from these and previous results (10, 11, 13, 22) in which inefficient negative selection results in the escape of a self-reactive biased repertoire (marked by Vβ’s responsive to MMTV superantigens) that is further enriched in the periphery through chronic antigen encounter. Although naïve SKG CD4 T cells can upregulate inhibitory pathways and an anergic-like transcriptional program in response to chronic antigen encounter (e.g., *Nr4a1*, *Pdcd1*, *Lag3*), due to weak proximal TCR signaling they are unable to efficiently and fully upregulate these programs (e.g., lower expression of *Izumo1r*, *Tnfrsf9*, *Tigit*, *Tox*) resulting in a defective anergy program and reduced numbers of peripheral anergic cells (11). Furthermore, Treg peripheral tolerance mechanisms are also severely impaired due to their attenuated TCR signaling and their altered Treg repertoire (12), releasing another checkpoint on arthritogenic T cells. We propose that costimulatory molecules such as IL-6, and perhaps other cytokines, promote T cell survival and lower the TCR signaling threshold required for peripheral activation and differentiation thereby allowing the activation of naïve SKG T cells that failed to upregulate TCR signaling-induced tolerogenic programs as we observed on our trajectory analysis (65–67). Therefore, the biased TCR Vβ repertoire unique to the SKGNur GFP^hi^ CD4 T cells are transcriptionally on the brink of activation such that an innate immune stimulus can trigger these cells to become potential initiators, or amplifiers, of disease. Future studies will determine whether these Vβs are sufficient, and/or necessary, to cause or exacerbate SKG arthritis and the role of MMTV superantigen(s) in SKG arthritis development. These findings have relevant implications in human autoimmune disease where endogenous or foreign antigens can also prime ‘dormant’ autoreactive T cells and trigger disease.

## Materials and Methods

### Materials

#### Antibodies and reagents

Ghost Dye Violet 510 (Tonbo Biosciences: 13-0870-T100) was used for live/dead staining. The following antibodies were used for staining as indicated: CD3e-BUV395 (BD Bioscience: 563565, clone: 145-2C11), CD4-APCeFluor 780 (eBioscience: 47-0042-82, clone: RM4-5), CD25-PerCPCy5.5 (Tonbo BioL 65-0251-U100, clone: PC61.5), CD44-PE-Cy7 (BioLegend: 103030, clone: IM7), CD62L-BV711 (BioLegend: 104445, clone: MEL-14), TCR Vβ3-PE (BD Bioscience: 553209, clone: KJ25), TCR Vβ5.1/5.2-PE (BD Bioscience: 562088, clone: MR9-4), TCR Vβ6-BV421 (BD Bioscience: 744590, clone: RR4-7), TCR Vβ8-BV421 (BD Bioscience: 742376, clone: F23), TCR Vβ11-PE (BD Bioscience: 553198, clone: RR3-15), TCR Vβ14-Biotin (BD Bioscience: 553257, clone: 14-2), Streptavidin-BV421 (BioLegend: 405226), FOXP3-eFluor 660 (eBioscience: 50-5773-82, clone: FJK-16s).

### Methods

#### Mice

BALB/c and C57BL/6J mice were purchased from Jackson laboratory, and BALB/cNur77-eGFP and SKGNur77-eGFP mice were bred in our facility (University of California, San Francisco) as previously described (11). All mice were housed and bred in specific pathogen-free conditions in the Animal Barrier Facility at UCSF according to the University Animal Care Committee and NIH guidelines. All animal experiments were approved by the UCSF Institutional Animal Care and Use Committee.

#### Flow cytometry and cell sorting

Cells were stained with antibodies of the indicated specificities and analyzed on a BD LSR Fortessa flow cytometer. Flow cytometry plots and analyses were performed using FlowJo v.10.8.0 (Tree Star). Cells were sorted to >95% purity using a MoFlo XDP (Beckman Coulter).

#### Statistics

Data were analyzed by comparison of means using paired or unpaired 2-tailed Student’s *t* tests using Prism v.9.2.0 for Mac (GraphPad Software). Data in all figures represent mean ± SEM unless otherwise indicated. Differences were considered significant at *P <* 0.05: \**P* < 0.05, \*\**P <* 0.01, \*\*\**P <* 0.001, and \*\*\*\**P <* 0.0001.

#### Murine synovial tissue preparation

Synovial tissues from ankle joints were digested with 1 mg/mL Collagenase IV (Worthington: LS004188) and DNase I (Sigma: 4536282001) in RPMI 1640 medium for 2 h at 37 °C on a rotator then quenched with 10% fetal bovine serum in RPMI 1640 medium; digested cells were filtrated through a 70 μm nylon mesh to prepare single cell suspensions.

#### Surface and intracellular staining

After live/dead staining with Ghost Dye Violet 510 as per manufacturer’s instructions, cells were stained for surface markers, washed, and then fixed for 10 min with 4% (vol/vol) fresh paraformaldehyde at room temperature protected from light. Cells were then permeabilized using the Mouse Regulatory T-Cell Staining kit 1 (eBioscience: 00-5521-00) per manufacturer’s instruction and then stained with FoxP3 e660.

#### *In vivo* treatments

Zymosan A (Sigma-Aldrich) suspended in saline at 10 mg/mL was kept in boiling water for 10 min. Zymosan A solution 2 mg or saline was intraperitoneally injected into 8-12 week-old mice.

#### PCR and RT-PCR

BALB/cJ, C57BL/6J, and SKG tail DNA was typed for *Mtv-6, -8, -9,* and *-17*. Standard PCR protocols were used for preparing PCR mixtures. Primer pairs for the detection of MMTV proviruses were previously described (68). GAPDH primers used: (5′ CATGTTTGTGATGGGTGTGAACCA 3′) and (5′ GTTGCTGTAGCCGTATTCATTGTC 3′). PCR mixtures for *Mtv-6, -8, and -9* were incubated at 94°C for 5 min, then denatured for 44 cycles at 94°C for 1 min, annealed at 46°C for 1 min, polymerized at 72°C for 1 min, and then incubated at 72°C for 5 min. PCRs for *Mtv-17* were conducted similarly except for an annealing temperature of 50°C. Samples were run on 2% agarose gel.

##### RT-PCR with joints

Single cells suspensions of synovial tissues from SKG ankle joints were spun down at 1500RPM at 4C. Cell pellets were flash frozen using dry ice in ethyl alcohol. Frozen cell pellets were used with the RNeasy Mini Kit (Qiagen: 74106) for RNA purification. The qScript cDNA Synthesis Kit (Quantabio: 95047-100) was used for cDNA library synthesis from purified total RNA. RT-PCR was conducted as described previously for PCR.

#### Bulk RNA sequencing

Negatively selected CD4 T cells from the lymph node were sorted for CD62L^hi^CD44^lo^CD25^-^ and the 10% highest (GFP^hi^) or lowest (GFP^lo^) expressing T cells. Cells were washed, pelleted and immediately flash frozen using dry ice in ethyl alcohol. Samples were processed for bulk RNA-sequencing by Q2 solutions using the TruSeq Stranded mRNA kit (Illumina: RS-122-2103) for library preparation. The resulting libraries pool into three batches and sequenced on a Illumina HiSeq 2500 sequencer over three lanes.

#### Alignment and initial processing of bulk RNA sequencing data

The raw fastq files were clipped and filtered using fastq-mcf v.1.04.636 to remove low quality reads and bases, homopolymers, and adapter sequences. The filtered reads were aligned using the STAR v.2.4 (69) with the default settings to the mm10 transcriptome to produce count matrices for each sample. Genes with less than 10 counts across all the samples were filtered out. Raw counts were normalized and transformed by the variance stabilizing transformation (VST) function from DESeq2 v.1.22.2 (70).

#### PCA analysis

The VST normalized features were used for principal component analysis with the function plotPCA from DESeq2.

#### Bulk RNA sequencing differential expression

Differential gene expression for the bulk RNA sequencing samples was performed with the raw counts from the filtered gene list for the indicated samples as the inputs. The analysis was run using a negative binomial model with multiple testing correction with Benjamini-Hochberg implemented via the DESeq function which includes an internal normalization from DESeq2. For differential gene expression between samples within the same genotype, mouse identity was included as a covariate.

#### Functional enrichment analysis

The collection of 991 significantly differentially expressed genes (log2FC > 1 and adjusted p value < 0.05) from the four comparisons [SKGNur GFP^hi^ versus SKGNur GFP^lo^, WTNur GFP^hi^ versus WTNur GFP^lo^, SKGNur GFP^hi^ versus WTNur GFP^hi^, SKGNur GFP^lo^ versus WTNur GFP^lo^] were hierarchically clustered using the Ward linkage (“ward.D2”) with the R package pheatmap v.1.0.12. The resulting dendrogram was used to partition the differentially expressed gene list into six gene modules. The gene lists for each gene module were analyzed using the functional profiling g:GOSt tool from g:Profiler (version e102_eg49_p15_e7ff1c9) with g:SCS multiple testing correction method applying significance threshold of 0.05. Select significantly enriched pathways from the GO:BP or KEGG collections were reported.

#### Gene set enrichment analysis

For the bulk RNA differential expression, the differential gene list was filtered to remove genes with NA for the adjusted p value or log fold change. For the single-cell RNA differential expression, the differential gene list was filtered to only include genes which were expressed in at least 1% of cells in the T.4N*_Nr4a1_* cluster. These filtered gene list were used to create ranked gene lists with the sign(log fold change) times the −log10(raw p value) as the ranking metric. The ranked list was used as input to look for gene set enrichment in the indicated collection of pathways in the ‘classic’ mode with the GSEAPreranked tool from GSEA v.4.1.0 with the default settings. For pathway collections of human genes, the ‘Mouse_Gene_Symbol_Remapping_Human_Orthologs_MSigDB’ chip file was used to map mouse genes from the ranked gene list to the human orthologs. Mouse gene symbols that mapped to the same human symbol were collapsed based on the max rank.

#### Single-cell RNA and TCR sequencing

Negatively selected CD4 T cells from the lymph node and spleen were sorted for CD62L^hi^CD44^lo^CD25^-^ and the 10% highest (GFP^hi^) or lowest (GFP^lo^) expressing T cells. Droplet-based paired single-cell RNA and TCR sequencing was performed using the 10x single-cell 5’+V(D)J v.1 kit per manufacterer’s instructions. The resulting cDNA libraries were sequenced on four lanes of an Illumina Novaseq 6000 sequencer to yield gene expression (GEX) and T cell receptor (TCR) fastqs.

#### Alignment and initial processing of single-cell sequencing data

The raw fastq files were aligned using CellRanger v3.0.1 and 3.0.2 software with the default settings to the mm10 transcriptome with the addition of the sequence for the eGFP transcript and the vdj GRCm38 v 3.1.0 reference for the GEX and TCR fastqs, respectively.

#### eGFP transcript sequence

ATGGTGAGCAAGGGCGAGGAGCTGTTCACCGGGGTGGTGCCCATCCTGGTCGAGCTGGACGGCGACGTAAACGGCCACAAGTTCAGCGTGTCCGGCGAGGGCGAGGGCGATGCCACCTACGGCAAGCTGACCCTGAAGTTCATCTGCACCACCGGCAAGCTGCCCGTGCCCTGGCCCACCCTCGTGACCACCCTGACCTACGGCGTGCAGTGCTTCAGCCGCTACCCCGACCACATGAAGCAGCACGACTTCTTCAAGTCCGCCATGCCCGAAGGCTACGTCCAGGAGCGCACCATCTTCTTCAAGGACGACGGCAACTACAAGACCCGCGCCGAGGTGAAGTTCGAGGGCGACACCCTGGTGAACCGCATCGAGCTGAAGGGCATCGACTTCAAGGAGGACGGCAACATCCTGGGGCACAAGCTGGAGTACAACTACAACAGCCACAACGTCTATATCATGGCCGACAAGCAGAAGAACGGCATCAAGGTGAACTTCAAGATCCGCCACAACATCGAGGACGGCAGCGTGCAGCTCGCCGACCACTACCAGCAGAACACCCCCATCGGCGACGGCCCCGTGCTGCTGCCCGACA ACCACTACCTGAGCACCCAGTCCGCCCTGAGCAAAGACCCCAACGAGAAGCGCGATCACATGGTCCTGCTGGAGTTCGTGACCGCCGCCGGGATCACTCTCGGCATGGACGAGCTGTACA AGTAA

#### Cell type classification and clustering

We filtered out 721 cells with less than 100 or more than 3000 genes detected and filtered out 14,388 genes detected in less than 3 cells. We also filtered out 1,066 cells with more than 10% of total counts (UMIs) mapping to mitochondrial genes and 1008 cells determined to be contaminating B cells based on CD19 expression. The raw counts were normalized to 10,000 counts per cell and log(count + 1) transformed. For technical and batch correction, we regressed out total UMI counts and % counts mapping to mitochondrial genes and used combat for batch correction with each sample as a batch. We identified 1119 highly variable genes (excluding all Trav and Trbv genes to avoid clustering cells based on expression of those genes) which were scaled and used with the default settings in scanpy v.1.4.3 (71) for PCA analysis followed by leiden clustering after nearest neighbor detection and UMAP projection. This analysis identified 13 clusters which we collapsed into 9 cell sub-types based on differential gene analysis.

#### Single-cell differential expression analysis

Differential gene expression on the log-normalized gene counts for the SKGNur GFP^hi^ cells versus the SKGNur GFP^lo^ cells with a particular TRBV assignment (**Fig. 5G**, **fig. S4D**) was performed using the hurdle model with Benjamini-Hochberg multiple testing correction from MAST v.1.12.0 (72) to allow for paired differential expression by mouse with the formula: ~ genotype + cell sub-type + number of genes detected + mouse id. All other single-cell differential expression was performed using the log-normalized gene counts with the rank_genes_groups function from scanpy with the Wilcoxon rank-sum method and multiple testing correction with Benjamini-Hochberg. Additionally, the adjusted p values that were equal to 0 were updated to the minimum representable positive normalized float (2.2250738585072014e-308).

#### Cell cycle phase assignment and module scoring

To assign cells to the cell cycle phases, the log-normalized scaled gene counts were used with the score_genes_cell_cycle function from the scanpy v.1.5.1 package with the *Mus musculus* G1/S DNA Damage Checkpoints and G2/M Checkpoints gene lists from the REACTOME database being used for the genes associated to the S phase and genes associated to the G2M phase (73, 74), respectively. For the single cell scoring of the bulk RNA sequencing gene modules, the log-normalized scaled gene counts were used with the score_genes function from scanpy.

#### RNA velocity analysis

For each 10x well, we used velocyto v.0.17.17 (75) to create a loom file with the spliced, unspliced, and ambiguous counts with the Dec. 2011 GRCm38/mm10 repeat masking gtf file from the UCSC genome browser (76, 77). The loom files across all wells were merged and then subsetted to all cells in the T.4N*_Nr4a1_* cluster. The resulting object was used to run the dynamical model from scvelo v.0.2.1 with the default settings to uncover the RNA velocity to predict the latent time for each cell.

We used we used a Gaussian mixture model with the GaussianMixture tool from sklearn v.0.23.1 (78) to deconvolute the underlying individual Gaussian distributions from the latent time distribution for cells from the T.4N*_Nr4a1_* cluster. This separated the cells into an optimal number of 4 distributions or clusters as determined by the elbow of the Bayesian Information Criterion (BIC) and Akaike Information Criterion (AIC) plots.

The smoothed gene expression versus latent time was modelled using a linear generalized additive model using default settings with the LinearGAM function from pygam v.0.8.0 (79). For trajectory inference between the four clusters (“Stage 1” – “Stage 4”), we used the graph-based tool PAGA within scvelo to predict velocity-inferred transitions among the clusters.

#### TCR analysis

Cells with <=2 TRA chains and <=1 TRB chains were used in the TCR clonotype analyses (53). Cells with two TRA chains were removed for the TRBV and TRAV analyses since the highest frequency for any dual TRA was 0.09% in any one sample (~1 cell). This removed 10,598 cells or 13.6% of all cells which is consistent with the expected dual TRA frequency. TRBV and TRAV genes which were not present in at least two mice from the same subgroup (i.e., SKGNur GFP^hi^, WTNur GFP^hi^, SKGNur GFP^lo^, and WTNur GFP^lo^) were removed from the downstream TRBV and TRAV analyses.

Significant differences in the TRBV frequencies between subgroups was determined by exact permutation test for unpaired samples and exact permutation test (N > 5 paired samples) (80) or paired t-test (N <= 5 paired samples) using scipy v.1.4.1 followed by Benjamini-Hochberg correction with statsmodels v.0.11.1 for paired samples.

## Acknowledgments

We thank Lewis Lanier, Julie Zikherman, and Philippa Marrack for valuable comments that improved the manuscript. We thank Z. Wang for cell sorting, A. Roque for animal husbandry, and Chan Zuckerberg Biohub for single-cell sequencing.

This research was supported by NIH grants K08AR072144 (to J.F.A.), T32GM007618 and F30CA257291 (to E.M.), R01HG011239, R01AI136972, and U01HG012192 (to C.J.Y.), R37AI114575 (to A.W.). A.W. is supported by Howard Hughes Medical Institute. J.D. and C.J.Y. are supported by the Chan Zuckerberg Biohub. C.J.Y. is also supported by the Chan Zuckerberg Initiative and the Parker Institute for Cancer Immunotherapy (PICI). J.F.A. is supported by the UCSF Center for the Rheumatic Diseases and the Rosalind Russell Medical Research Foundation Bechtel Award. Funding for this work was also supported by the Arthritis National Research Foundation (to J.F.A.) and the UCSF Research Evaluation Allocation Committee funded by Esther Memorial fund (to J.F.A).

## Author contributions

Conceptualization: J.F.A., A.W., C.J.Y., J.D.

Methodology: J.F.A, C.J.Y, A.W., E.M., S.Y.

Investigation: J.F.A, E.M., S.Y., C.L., N.P. Formal Analysis: J.F.A, E.M., C.J.Y. Visualization: J.F.A, E.M., S.Y., C.L., N.P.

Funding acquisition: J.F.A., A.W., C.J.Y., J.D.

Project administration: J.F.A., A.W., C.J.Y., J.D., E.M.

Resources: J.F.A., A.W., C.J.Y., J.D.

Supervision: J.F.A., A.W., C.J.Y. Writing – original draft: J.F.A, E.M.

Writing – review & editing: J.F.A., E.M., A.W., C.J.Y.

## Data and materials availability

Bulk RNA sequencing data and single-cell RNA and TCR sequencing data will be released on NCBI GEO under accession number GSE185577 upon manuscript acceptance. All other data are available in the main text or as supplementary data.

## Code availability

Code for analysis is available at: https://github.com/yelabucsf/SKG_rheum for review. It will be uploaded to Zenodo upon acceptance of the manuscript.

## Supplementary Materials

Supplementary Figures 1 to 6

Supplementary Data 1 to 10

## References

1. Cope AP, Schulze-Koops H, Aringer M. The central role of T cells in rheumatoid arthritis. Clinical and experimental rheumatology. 2007;25(5 Suppl 46):S4-11.

2. Plenge RM, Seielstad M, Padyukov L, Lee AT, Remmers EF, Ding B, et al. TRAF1-C5 as a risk locus for rheumatoid arthritis--a genomewide study. N Engl J Med. 2007;357(12):1199-209.

3. Schmidt D, Goronzy JJ, Weyand CM. CD4+ CD7-CD28-T cells are expanded in rheumatoid arthritis and are characterized by autoreactivity. The Journal of clinical investigation. 1996;97(9):2027-37.

4. Thomas R, McIlraith M, Davis LS, Lipsky PE. Rheumatoid synovium is enriched in CD45RBdim mature memory T cells that are potent helpers for B cell differentiation. Arthritis and rheumatism. 1992;35(12):1455-65.

5. Weyand CM, Fujii H, Shao L, Goronzy JJ. Rejuvenating the immune system in rheumatoid arthritis. Nature reviews Rheumatology. 2009;5(10):583-8.

6. Zhang Z, Gorman CL, Vermi AC, Monaco C, Foey A, Owen S, et al. TCRzetadim lymphocytes define populations of circulating effector cells that migrate to inflamed tissues. Blood. 2007;109(10):4328-35.

7. Gringhuis SI, Papendrecht-van der Voort EA, Leow A, Nivine Levarht EW, Breedveld FC, Verweij CL. Effect of redox balance alterations on cellular localization of LAT and downstream T-cell receptor signaling pathways. Molecular and cellular biology. 2002;22(2):400-11.

8. Maurice MM, Lankester AC, Bezemer AC, Geertsma MF, Tak PP, Breedveld FC, et al. Defective TCR-mediated signaling in synovial T cells in rheumatoid arthritis. Journal of immunology (Baltimore, Md: 1950). 1997;159(6):2973-8.

9. Romagnoli P, Strahan D, Pelosi M, Cantagrel A, van Meerwijk JP. A potential role for protein tyrosine kinase p56(lck) in rheumatoid arthritis synovial fluid T lymphocyte hyporesponsiveness. International immunology. 2001;13(3):305-12.

10. Sakaguchi N, Takahashi T, Hata H, Nomura T, Tagami T, Yamazaki S, et al. Altered thymic T-cell selection due to a mutation of the ZAP-70 gene causes autoimmune arthritis in mice. Nature. 2003;426(6965):454-60.

11. Ashouri JF, Hsu L-Y, Yu S, Rychkov D, Chen Y, Cheng DA, et al. Reporters of TCR signaling identify arthritogenic T cells in murine and human autoimmune arthritis. Proceedings of the National Academy of Sciences. 2019;116(37):18517-27.

12. Tanaka S, Maeda S, Hashimoto M, Fujimori C, Ito Y, Teradaira S, et al. Graded attenuation of TCR signaling elicits distinct autoimmune diseases by altering thymic T cell selection and regulatory T cell function. Journal of immunology (Baltimore, Md: 1950). 2010;185(4):2295-305.

13. Hsu LY, Tan YX, Xiao Z, Malissen M, Weiss A. A hypomorphic allele of ZAP-70 reveals a distinct thymic threshold for autoimmune disease versus autoimmune reactivity. J Exp Med. 2009;206(11):2527-41.

14. Yoshitomi H, Sakaguchi N, Kobayashi K, Brown GD, Tagami T, Sakihama T, et al. A role for fungal {beta}-glucans and their receptor Dectin-1 in the induction of autoimmune arthritis in genetically susceptible mice. J Exp Med. 2005;201(6):949-60.

15. Sakaguchi S, Benham H, Cope AP, Thomas R. T-cell receptor signaling and the pathogenesis of autoimmune arthritis: insights from mouse and man. Immunology and cell biology. 2012;90(3):277-87.

16. Evans RM. The steroid and thyroid hormone receptor superfamily. Science (New York, NY). 1988;240(4854):889-95.

17. Olefsky JM. Nuclear receptor minireview series. The Journal of biological chemistry. 2001;276(40):36863-4.

18. Deane KD, Holers VM. The Natural History of Rheumatoid Arthritis. Clin Ther. 2019;41(7):1256-69.

19. Raudvere U, Kolberg L, Kuzmin I, Arak T, Adler P, Peterson H, et al. g:Profiler: a web server for functional enrichment analysis and conversions of gene lists (2019 update). Nucleic acids research. 2019;47(W1):W191-w8.

20. Mootha VK, Lindgren CM, Eriksson KF, Subramanian A, Sihag S, Lehar J, et al. PGC-1alpha-responsive genes involved in oxidative phosphorylation are coordinately downregulated in human diabetes. Nature genetics. 2003;34(3):267-73.

21. Subramanian A, Tamayo P, Mootha VK, Mukherjee S, Ebert BL, Gillette MA, et al. Gene set enrichment analysis: a knowledge-based approach for interpreting genome-wide expression profiles. Proc Natl Acad Sci U S A. 2005;102(43):15545-50.

22. Ito Y, Hashimoto M, Hirota K, Ohkura N, Morikawa H, Nishikawa H, et al. Detection of T cell responses to a ubiquitous cellular protein in autoimmune disease. Science (New York, NY). 2014;346(6207):363-8.

23. Myers DR, Lau T, Markegard E, Lim HW, Kasler H, Zhu M, et al. Tonic LAT-HDAC7 Signals Sustain Nur77 and Irf4 Expression to Tune Naive CD4 T Cells. Cell reports. 2017;19(8):1558-71.

24. Myers DR, Zikherman J, Roose JP. Tonic Signals: Why Do Lymphocytes Bother? Trends in immunology. 2017;38(11):844-57.

25. Nguyen TTT, Wang ZE, Shen L, Schroeder A, Eckalbar W, Weiss A. Cbl-b deficiency prevents functional but not phenotypic T cell anergy. J Exp Med. 2021;218(7).

26. Ashouri JF, Weiss A. Endogenous Nur77 Is a Specific Indicator of Antigen Receptor Signaling in Human T and B Cells. Journal of immunology (Baltimore, Md: 1950). 2017;198(2):657-68.

27. Huang B, Pei HZ, Chang HW, Baek SH. The E3 ubiquitin ligase Trim13 regulates Nur77 stability via casein kinase 2alpha. Scientific reports. 2018;8(1):13895.

28. Tan C, Hiwa R, Mueller JL, Vykunta V, Hibiya K, Noviski M, et al. NR4A nuclear receptors restrain B cell responses to antigen when second signals are absent or limiting. Nature immunology. 2020;21(10):1267-79.

29. Zhang L, Xie F, Zhang J, Dijke PT, Zhou F. SUMO-triggered ubiquitination of NR4A1 controls macrophage cell death. Cell Death Differ. 2017;24(9):1530-9.

30. ElTanbouly MA, Zhao Y, Nowak E, Li J, Schaafsma E, Le Mercier I, et al. VISTA is a checkpoint regulator for naïve T cell quiescence and peripheral tolerance. Science (New York, NY). 2020;367(6475).

31. Jennings E, Elliot TAE, Thawait N, Kanabar S, Yam-Puc JC, Ono M, et al. Nr4a1 and Nr4a3 Reporter Mice Are Differentially Sensitive to T Cell Receptor Signal Strength and Duration. Cell reports. 2020;33(5):108328.

32. Bour-Jordan H, Esensten JH, Martinez-Llordella M, Penaranda C, Stumpf M, Bluestone JA. Intrinsic and extrinsic control of peripheral T-cell tolerance by costimulatory molecules of the CD28/lf1B7 family. Immunological reviews. 2011;241(1):180-205.

33. Chen L, Flies DB. Molecular mechanisms of T cell co-stimulation and co-inhibition. Nature reviews Immunology. 2013;13(4):227-42.

34. Fife BT, Bluestone JA. Control of peripheral T-cell tolerance and autoimmunity via the CTLA-4 and PD-1 pathways. Immunological reviews. 2008;224:166-82.

35. Lee J, Su EW, Zhu C, Hainline S, Phuah J, Moroco JA, et al. Phosphotyrosine-dependent coupling of Tim-3 to T-cell receptor signaling pathways. Molecular and cellular biology. 2011;31(19):3963-74.

36. Kalekar LA, Schmiel SE, Nandiwada SL, Lam WY, Barsness LO, Zhang N, et al. CD4(+) T cell anergy prevents autoimmunity and generates regulatory T cell precursors. Nature immunology. 2016;17(3):304-14.

37. Mueller DL. E3 ubiquitin ligases as T cell anergy factors. Nature immunology. 2004;5(9):883-90.

38. Zheng Y, Zha Y, Driessens G, Locke F, Gajewski TF. Transcriptional regulator early growth response gene 2 (Egr2) is required for T cell anergy in vitro and in vivo. J Exp Med. 2012;209(12):2157-63.

39. Crawford A, Angelosanto JM, Kao C, Doering TA, Odorizzi PM, Barnett BE, et al. Molecular and transcriptional basis of CD4(+) T cell dysfunction during chronic infection. Immunity. 2014;40(2):289-302.

40. Trefzer A, Kadam P, Wang SH, Pennavaria S, Lober B, Akcabozan B, et al. Dynamic adoption of anergy by antigen-exhausted CD4(+) T cells. Cell reports. 2021;34(6):108748.

41. Chen J, Lopez-Moyado IF, Seo H, Lio CJ, Hempleman LJ, Sekiya T, et al. NR4A transcription factors limit CAR T cell function in solid tumours. Nature. 2019.

42. Liebmann M, Hucke S, Koch K, Eschborn M, Ghelman J, Chasan AI, et al. Nur77 serves as a molecular brake of the metabolic switch during T cell activation to restrict autoimmunity. Proc Natl Acad Sci U S A. 2018;115(34):E8017-e26.

43. Liu X, Wang Y, Lu H, Li J, Yan X, Xiao M, et al. Genome-wide analysis identifies NR4A1 as a key mediator of T cell dysfunction. Nature. 2019;567(7749):525-9.

44. Zinzow-Kramer WM, Weiss A, Au-Yeung BB. Adaptation by naïve CD4(+) T cells to self-antigen-dependent TCR signaling induces functional heterogeneity and tolerance. Proc Natl Acad Sci U S A. 2019;116(30):15160-9.

45. Ciofani M, Madar A, Galan C, Sellars M, Mace K, Pauli F, et al. A validated regulatory network for Th17 cell specification. Cell. 2012;151(2):289-303.

46. Yu CR, Mahdi RM, Ebong S, Vistica BP, Gery I, Egwuagu CE. Suppressor of cytokine signaling 3 regulates proliferation and activation of T-helper cells. The Journal of biological chemistry. 2003;278(32):29752-9.

47. Ye H, Zhang J, Wang J, Gao Y, Du Y, Li C, et al. CD4 T-cell transcriptome analysis reveals aberrant regulation of STAT3 and Wnt signaling pathways in rheumatoid arthritis: evidence from a case-control study. Arthritis research & therapy. 2015;17:76.

48. Atsumi T, Ishihara K, Kamimura D, Ikushima H, Ohtani T, Hirota S, et al. A point mutation of Tyr-759 in interleukin 6 family cytokine receptor subunit gp130 causes autoimmune arthritis. J Exp Med. 2002;196(7):979-90.

49. Shouda T, Yoshida T, Hanada T, Wakioka T, Oishi M, Miyoshi K, et al. Induction of the cytokine signal regulator SOCS3/CIS3 as a therapeutic strategy for treating inflammatory arthritis. The Journal of clinical investigation. 2001;108(12):1781-8.

50. Wong PK, Egan PJ, Croker BA, O’Donnell K, Sims NA, Drake S, et al. SOCS-3 negatively regulates innate and adaptive immune mechanisms in acute IL-1-dependent inflammatory arthritis. The Journal of clinical investigation. 2006;116(6):1571-81.

51. Bergen V, Lange M, Peidli S, Wolf FA, Theis FJ. Generalizing RNA velocity to transient cell states through dynamical modeling. Nature biotechnology. 2020;38(12):1408-14.

52. Wolf FA, Hamey FK, Plass M, Solana J, Dahlin JS, Göttgens B, et al. PAGA: graph abstraction reconciles clustering with trajectory inference through a topology preserving map of single cells. Genome biology. 2019;20(1):59.

53. Abdelnour A, Bremell T, Holmdahl R, Tarkowski A. Clonal expansion of T lymphocytes causes arthritis and mortality in mice infected with toxic shock syndrome toxin-1-producing staphylococci. European journal of immunology. 1994;24(5):1161-6.

54. Kappler J, Kotzin B, Herron L, Gelfand EW, Bigler RD, Boylston A, et al. V beta-specific stimulation of human T cells by staphylococcal toxins. Science (New York, NY). 1989;244(4906):811-3.

55. Hodes RJ, Abe R. Mouse endogenous superantigens: Ms and Mls-like determinants encoded by mouse retroviruses. Current protocols in immunology. 2001;Appendix 1:Appendix 1F.

56. Matsutani T, Ohmori T, Ogata M, Soga H, Yoshioka T, Suzuki R, et al. Alteration of T-cell receptor repertoires during thymic T-cell development. Scandinavian journal of immunology. 2006;64(1):53-60.

57. Günzburg WH, Heinemann F, Wintersperger S, Miethke T, Wagner H, Erfle V, et al. Endogenous superantigen expression controlled by a novel promoter in the MMTV long terminal repeat. Nature. 1993;364(6433):154-8.

58. Marrack P, Kushnir E, Kappler J. A maternally inherited superantigen encoded by a mammary tumour virus. Nature. 1991;349(6309):524-6.

59. Howell MD, Diveley JP, Lundeen KA, Esty A, Winters ST, Carlo DJ, et al. Limited T-cell receptor beta-chain heterogeneity among interleukin 2 receptor-positive synovial T cells suggests a role for superantigen in rheumatoid arthritis. Proc Natl Acad Sci U S A. 1991;88(23):10921-5.

60. Paliard X, West SG, Lafferty JA, Clements JR, Kappler JW, Marrack P, et al. Evidence for the effects of a superantigen in rheumatoid arthritis. Science (New York, NY). 1991;253(5017):325-9.

61. Jenkins RN, Nikaein A, Zimmermann A, Meek K, Lipsky PE. T cell receptor V beta gene bias in rheumatoid arthritis. The Journal of clinical investigation. 1993;92(6):2688-701.

62. Bhardwaj N, Hodtsev AS, Nisanian A, Kabak S, Friedman SM, Cole BC, et al. Human T-cell responses to Mycoplasma arthritidis-derived superantigen. Infection and immunity. 1994;62(1):135-44.

63. Grom AA, Thompson SD, Luyrink L, Passo M, Choi E, Glass DN. Dominant T-cell-receptor beta chain variable region V beta 14+ clones in juvenile rheumatoid arthritis. Proc Natl Acad Sci U S A. 1993;90(23):11104-8.

64. Zhao YX, Brunsberg U, Holmdahl R, Tarkowski A. V beta 11+ T-lymphocyte expansion by toxic shock syndrome toxin-1 differs in mice bearing H-2q versus H-2b haplotypes. Immunology. 1998;94(1):1-4.

65. Lotz M, Jirik F, Kabouridis P, Tsoukas C, Hirano T, Kishimoto T, et al. B cell stimulating factor 2/interleukin 6 is a costimulant for human thymocytes and T lymphocytes. J Exp Med. 1988;167(3):1253-8.

66. Rochman I, Paul WE, Ben-Sasson SZ. IL-6 increases primed cell expansion and survival. Journal of immunology (Baltimore, Md: 1950). 2005;174(8):4761-7.

67. Holsti MA, McArthur J, Allison JP, Raulet DH. Role of IL-6, IL-1, and CD28 signaling in responses of mouse CD4+ T cells to immobilized anti-TCR monoclonal antibody. Journal of immunology (Baltimore, Md: 1950). 1994;152(4):1618-28.

68. Barnett A, Mustafa F, Wrona TJ, Lozano M, Dudley JP. Expression of mouse mammary tumor virus superantigen mRNA in the thymus correlates with kinetics of self-reactive T-cell loss. Journal of virology. 1999;73(8):6634-45.

69. Dobin A, Davis CA, Schlesinger F, Drenkow J, Zaleski C, Jha S, et al. STAR: ultrafast universal RNA-seq aligner. Bioinformatics (Oxford, England). 2013;29(1):15-21.

70. Love MI, Huber W, Anders S. Moderated estimation of fold change and dispersion for RNA-seq data with DESeq2. Genome biology. 2014;15(12):550.

71. Wolf FA, Angerer P, Theis FJ. SCANPY: large-scale single-cell gene expression data analysis. Genome biology. 2018;19(1):15.

72. Finak G, McDavid A, Yajima M, Deng J, Gersuk V, Shalek AK, et al. MAST: a flexible statistical framework for assessing transcriptional changes and characterizing heterogeneity in single-cell RNA sequencing data. Genome biology. 2015;16:278.

73. G1/S DNA Damage Checkpoints. Reactome, v.75, https://reactome.org/content/detail/R-MMU-69615 [accessed January 16, 2021]. [Internet]. 2021 [cited January 16, 2021]. Available from: https://reactome.org/content/detail/R-MMU-69615.

74. G2/M Checkpoints. Reactome, v.75, https://reactome.org/content/detail/R-MMU-69473 [accessed January 16, 2021]. [Internet]. 2021 [cited January 16, 2021]. Available from: https://reactome.org/content/detail/R-MMU-69473.

75. La Manno G, Soldatov R, Zeisel A, Braun E, Hochgerner H, Petukhov V, et al. RNA velocity of single cells. Nature. 2018;560(7719):494-8.

76. Karolchik D, Hinrichs AS, Furey TS, Roskin KM, Sugnet CW, Haussler D, et al. The UCSC Table Browser data retrieval tool. Nucleic acids research. 2004;32(Database issue):D493-6.

77. University of California Santa Cruz. Genome Browser 2000 [updated July 20, 2021. Available from: http://genome.ucsc.edu

78. Pedregosa et al. Scikit-learn: Machine Learning in Python. JMLR 12, pp. 2825-2830 2011 [Available from: https://jmlr.csail.mit.edu/papers/v12/pedregosa11a.html.

79. Servén D. BC. dswah/pyGAM: v0.8.0. Zenodo. 2018 [Available from: https://doi.org/10.5281/zenodo.1476122.

80. Thompson WL, White GC, Gowan C. Detection of a Trend in Population Estimates. In: Thompson WL, White GC, Gowan C, editors. Monitoring Vertebrate Populations: Academic Press; 1998. p. 145-69.

