## Supplementary Figures 1-6 for "Naïve arthritogenic SKG T cells have a defect in anergy and a repertoire pruned by superantigen"

**This file includes:**

Supplementary Figs. 1 to 6

Captions for Data S1 to S10

**Other Supplementary Materials for this manuscript include the following:**

Data S1 to S10 [data_S1_bulk_RNA_seq_diff_exp.xlsx,

data_S2_heatmap_gene_list_with_modules.csv,

data_S3_GSEA_reports.xls,

data_S4_scRNAseq_diff_genes_by_cluster.csv,

data_S5_fig_2_diff_gene_lists.xlsx,

data_S6_diff_exp_Tnfrsf9_pos_Egr2_pos.csv,

data_S7_top_300_heatmap_gene_list.tsv,

data_S8_gini_coefficients.csv,

data_S9_TRBV_paired_test_nr4a1_cluster.csv,

data_s10_diff_exp_SKG_High_v_SKG_Low_for_TRBV.xlsx]

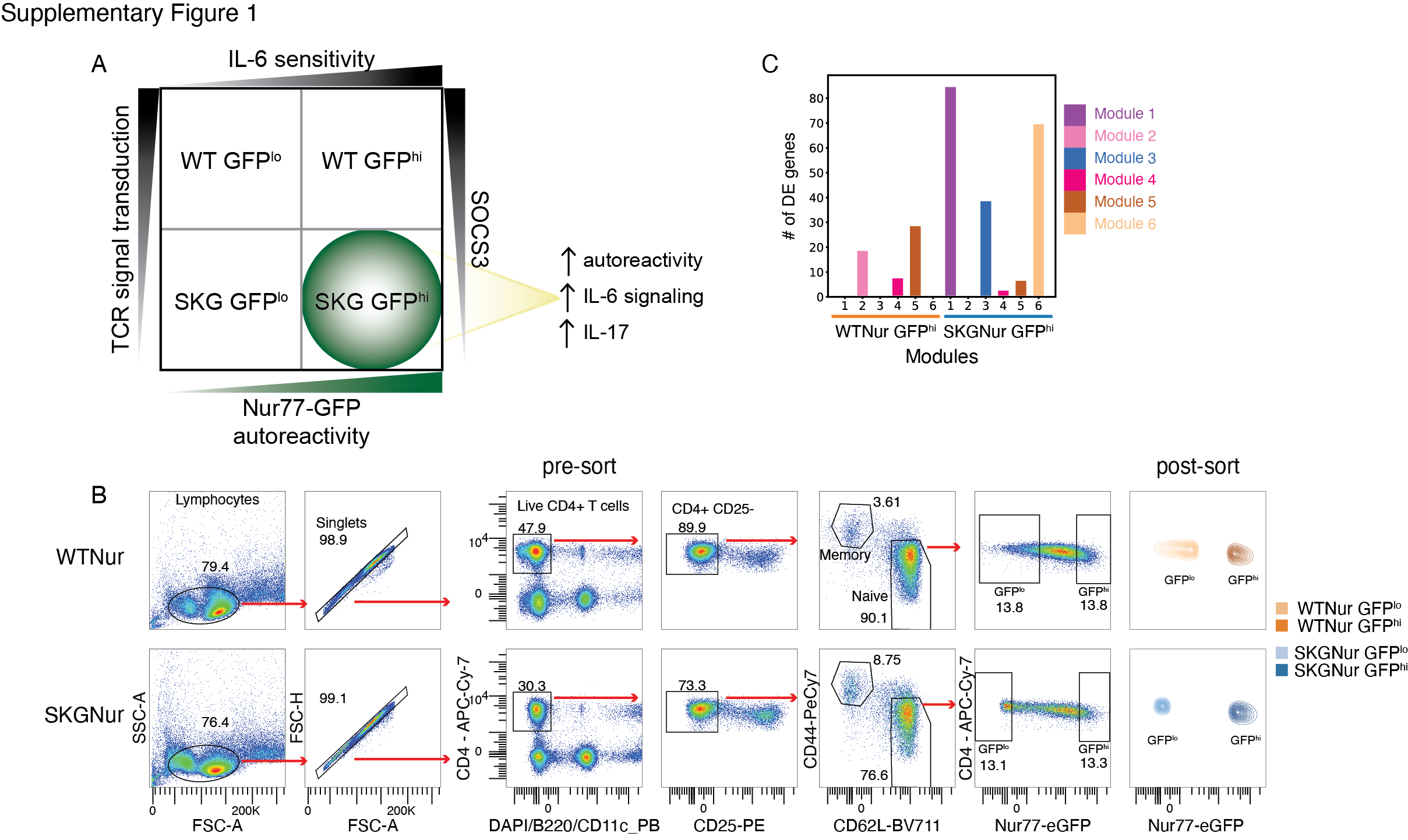

**Supplementary Fig. 1.** **SKGNur GFP^hi^ CD4 T cells readily differentiate into pathogenic effector cells.** (**A**) 2 x 2 matrix demonstrates how impaired TCR signaling observed in SKG mice (left y-axis, due to the hypomorphic Zap70 allele), in addition to chronic antigen stimulation (x-axis, resulting in higher levels of Nur77-eGFP demarcated by GFP^hi^) confer heightened sensitivity to IL-6 cytokine signaling, in part due to decreased levels of SOCS3. This contributes to the increased arthritogenicity observed in the autoreactive T cell clones that more readily differentiate into IL-17 producing CD4 T cells in SKG mice. (**B**) Gating for bulk RNAseq sorting of WTNur and SKGNur lymphocytes. (**C**) Bar plot of number of DEGs from WTNur GFP^hi^ and SKGNur GFP^hi^ cells contained in each gene module from **Fig. 1C**.

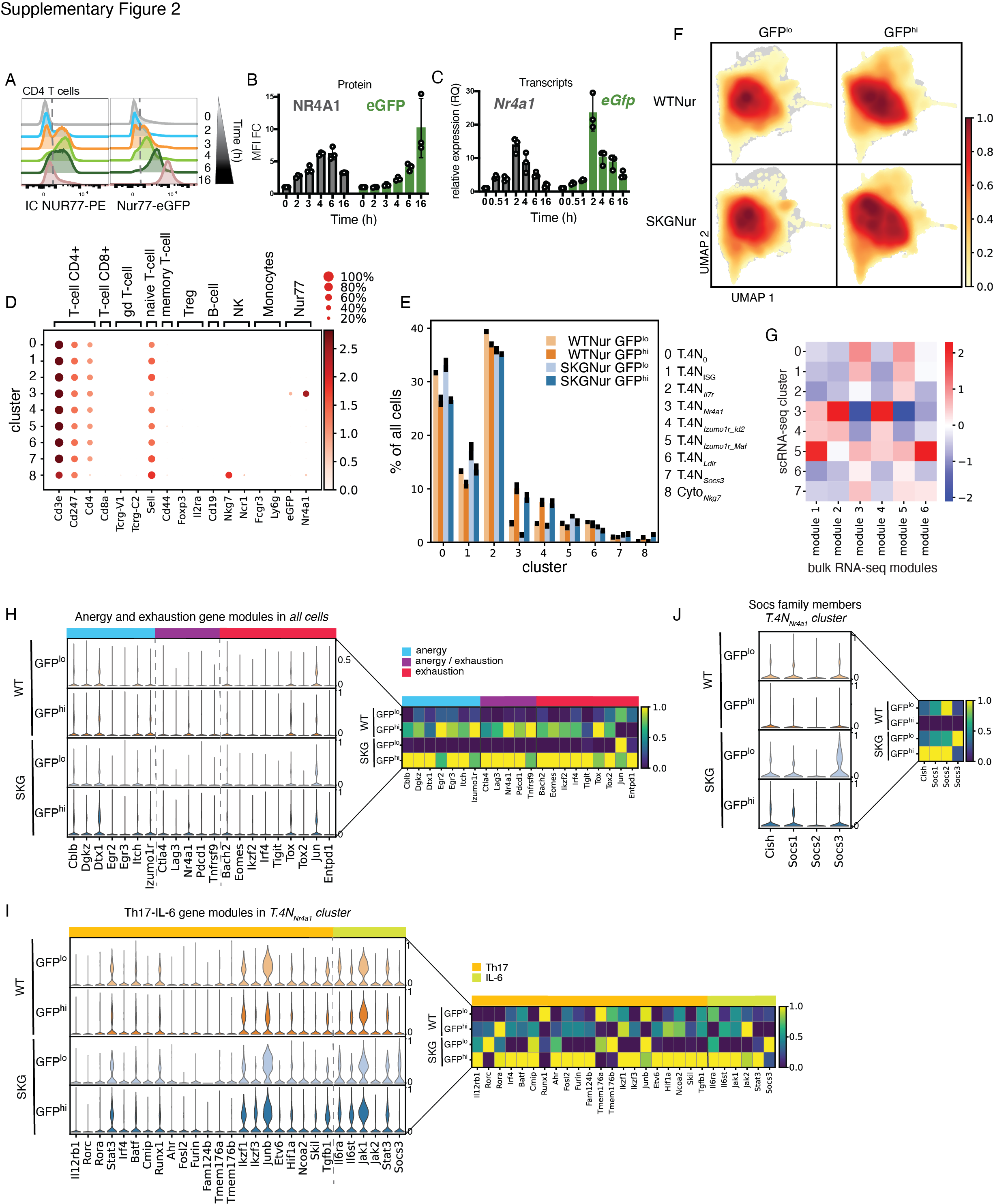

**Supplementary Fig. 2. NUR77/*Nr4a1* identifies naïve CD4 T cells that have recently encountered endogenous antigen resulting in a unique transcriptional program.** (**A-C**) CD4 T cells were stimulated ± platebound aCD3e + CD28 at indicated times. Levels of NUR77 and eGFP are depicted in representative histogram from 2 experiments (**A**) and MFI fold change (FC) quantified in (**B**) from 3 biological replicates. (**C**) Real-time RT-PCR measuring *Nr4a1* and *eGfp* mRNA levels in stimulated CD4 T cells from 3 biological replicates, from 2 independent experiments. (**D**) Expression of labelled genes for each cluster is shown by percentage of cells with expression greater than zero (dot size) and mean expression (color). (**E**) Bar plot of mean frequency for each subgroup of cells within each cluster. Black bars indicate difference between mouse 1 and mouse 2 for each subgroup. (**F**) UMAP colored by density of cells in all four subgroups (each subgroup contains samples from 2 mice). (**G**) Heatmap normalized by standard scale (subtract minimum and divide by maximum) by column of average single cell gene set scores for each cluster (excluding cluster 8 – Cyto_Nkg7_) for the gene sets defined by the modules from Fig. 1C. (**H-J**) Stacked violin plot demonstrates expression of candidate anergy and exhaustion associated genes (**H**), Th-17 and IL-6 associated genes (**I**), and *Socs* family members (**J**) in WTNur and SKGNur GFP^lo^ and GFP^hi^ CD4 naïve T cells in all cells (**H**) or in T.4N*_Nr4a1_* cells (**I-J**). Heatmaps on the right for each panel highlights differential expression of the indicated genes across subgroup by showing mean gene expression normalized by standard scale for each column.

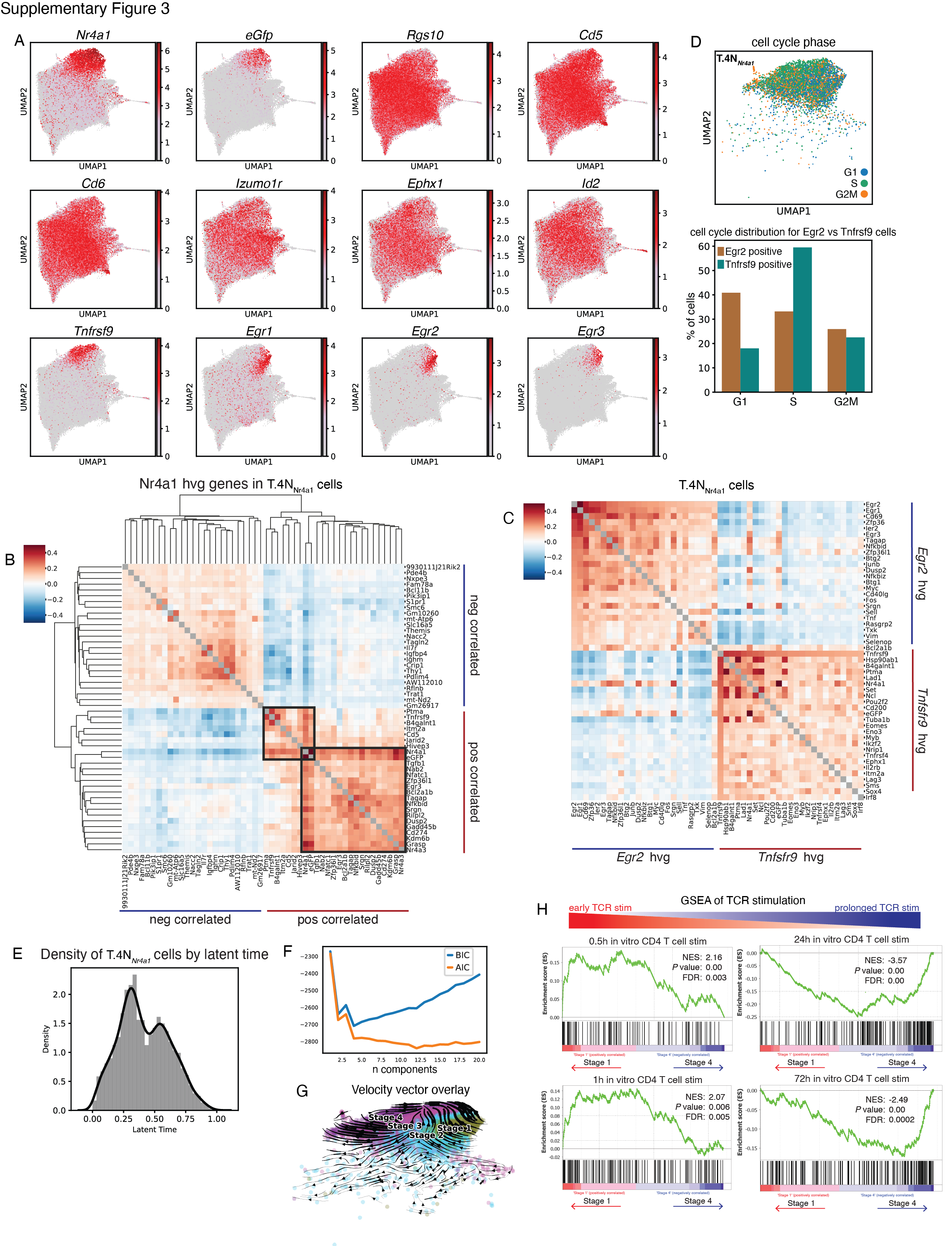

**Supplementary Fig. 3. Highly variable genes that positively and negatively correlate with *Nr4a1* in T.4N*_Nr4a1_* cluster and trajectory analysis of their underlying states.** (**A**) UMAP colored by expression levels of indicated genes from modules of genes positively correlating with *Nr4a1* as identified in Fig. 3A. Scale shows the log-normalized gene expression. (**B**) Hierarchical clustering of correlation matrix of top 25 HVGs that positively and negatively correlate with *Nr4a1* expression in T.4N*_Nr4a1_* cells using Spearman’s correlation. Diagonal grey colored boxes represent identity correlation of 1. Dark grey boxes mark modules of HVGs that highly correlate with *Nr4a1* expression. (**C**) Correlation matrix of HVGs that positively correlate with *Egr2* and *Tnfsrsf9* expression in T.4N*_Nr4a1_* cells using Spearman’s correlation. Diagonal grey boxes represent correlation of 1. (**D**) UMAP of cells from T.4N*_Nr4a1_* cluster colored by cell cycle phase assignment. Bar plot of % of cells in each cell cycle stage for cells expressing *Egr2* or *Tnfrsf9* (log-normalized expression > 1). (**E**) Probability density of latent time distribution of all cells in T.4N*_Nr4a1_* cluster. (**F**) Line plots for the Bayesian Information Criterion (BIC) and Akaike Information Criterion (AIC) for the Gaussian mixture model deconvolution versus number of underlying distributions or clusters. (**G**) UMAP colored by cell stage as defined in Fig. 3H with an overlay of velocity vectors for cell transitions as determined by the scvelo dynamical model. (**H**) Enrichment plots of pathways of time course *in vitro* activation of CD4+ T cells with aCD3 + CD28 from GSEA analysis of pathways from study GSE17974 for ranked genes from differential gene expression analysis of cells in Stage 1 versus Stage 4. FDR, false discovery rate. NES, normalized enrichment score.

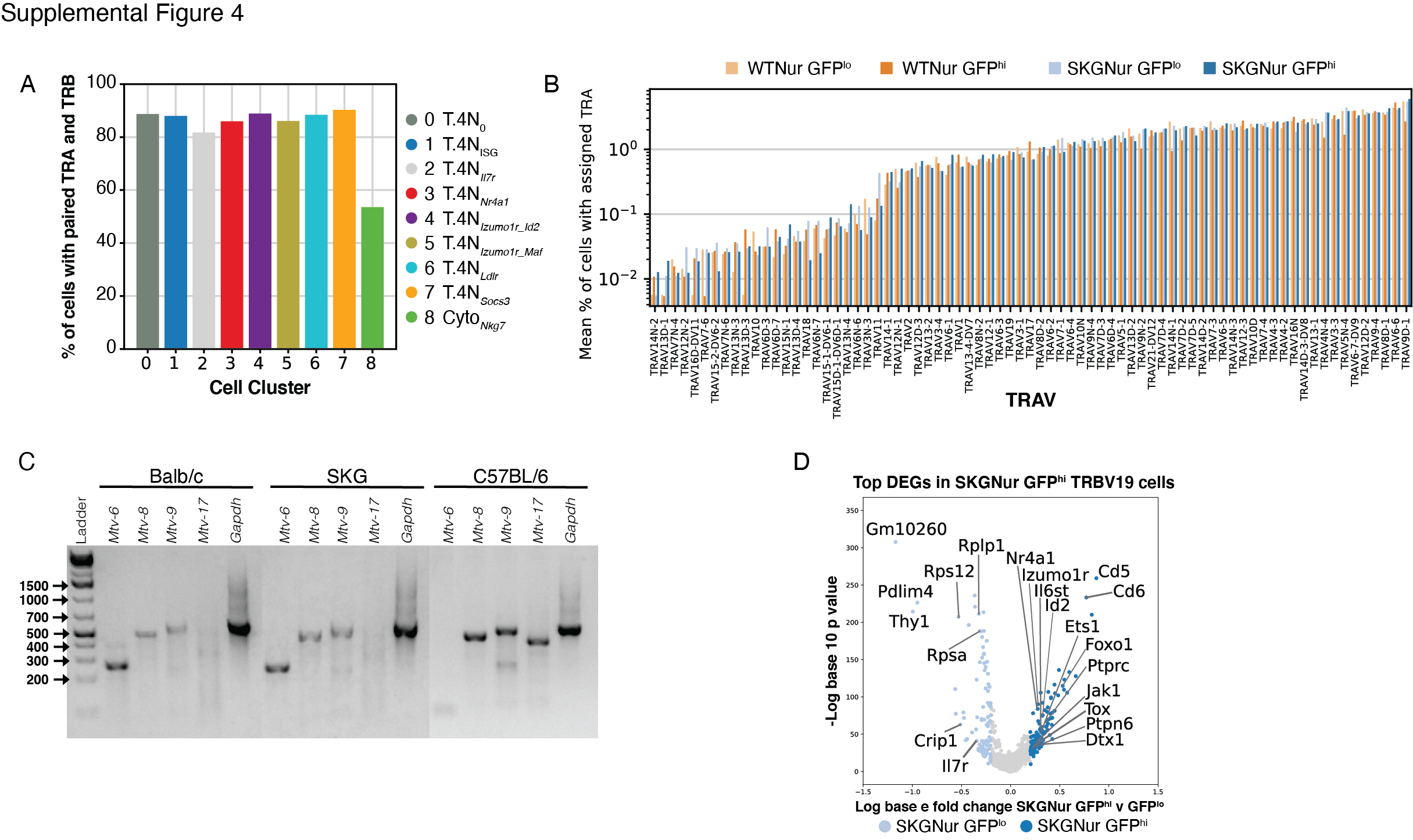

**Supplementary Fig. 4.** **SKGNur mice express superantigens involved in selection of TCR variable beta repertoire.** (**A**) Bar plot with percent of cells with paired TRA and TRB detection by cluster. (**B**) Bar plot of mean frequency of cells expressing each TRAV gene as a percentage of all cells in each sample with an assigned TRAV. Bars are colored according to subgroup and ordered by increasing overall frequency. (**C**) Balb/c and SKG tail DNA used in PCR reactions containing primers specific for the indicated Mtv pro-viruses. (**D**) Volcano plot of DEGs of cells with assigned indicated TRBV from the SKGNur GFP^hi^ or SKGNur GFP^lo^ subgroups. Dots are colored by significant overexpression (absolute value(natural log(fold-change)) > 0.2, adjusted *P value* < 0.05) in SKGNur GFP^hi^ (dark blue), SKGNur GFP^lo^ (light blue), or no significant difference (light grey).

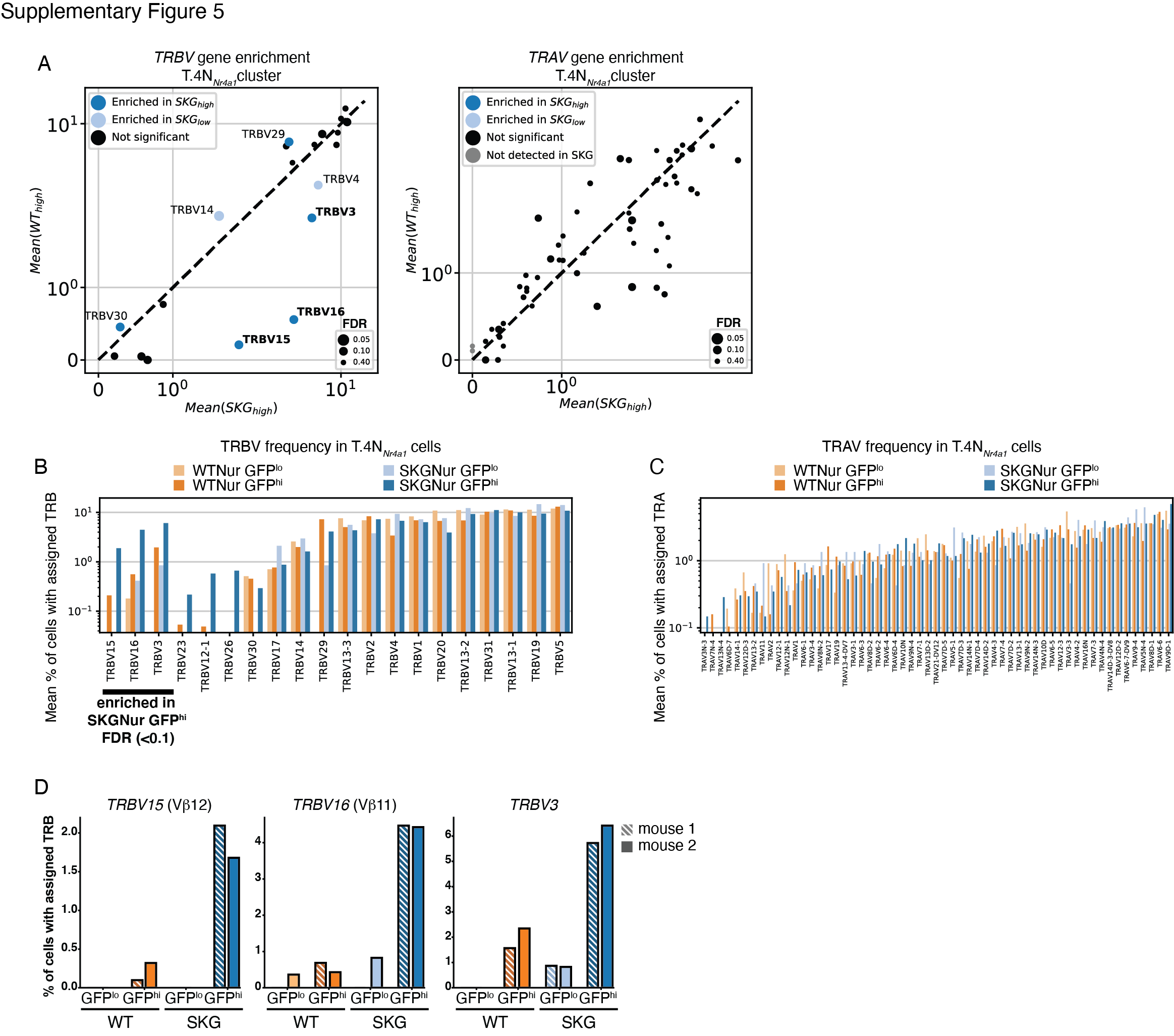

**Supplementary Fig. 5.** **Further enrichment of biased TRBV in SKGNur GFP^hi^ T.4N*_Nr4a1_* cells.** (**A**) Scatterplot of mean frequency of cells expressing each TRBV (left) or TRAV (right) gene for the SKGNur GFP^hi^ versus the WTNur GFP^hi^ T.4N*_Nr4a1_* cells. Dots for each TRV gene are sized according to the false discovery rate (FDR) from a paired one-sided t-test comparing frequency in SKGNur GFP^hi^ versus SKGNur GFP^lo^. Dots are colored as either significantly enriched (FDR < 0.1) in SKGNur GFP^hi^ (dark blue), significantly enriched in SKGNur GFP^lo^ (light blue), or not significantly enriched in either subgroup (black). TRBV genes that were significantly enriched in SKGNur GFP^hi^ and were also more highly expressed in SKGNur GFP^hi^ versus WTNur GFP^hi^ T.4N*_Nr4a1_* cells are bolded. (**B**) Bar plot of mean value of T.4N*_Nr4a1_* cells expressing each TRBV gene as a percentage of all T.4N*_Nr4a1_* cells in each sample with an assigned TRBV. Bars are colored according to subgroup and are ordered with the TRBV genes enriched in SKGNur GFP^hi^ T.4N*_Nr4a1_* cells (see fig. S5A) followed by the other TRBV genes ordered by increasing overall frequency. (**C**) Bar plot of mean value of T.4N*_Nr4a1_* cells expressing each TRAV gene as a percentage of all T.4N*_Nr4a1_* cells in each sample with an assigned TRAV. Bars are colored according to subgroup and are ordered by increasing overall frequency. (**D**) Bar plot of frequency of cells expressing indicated TRBV genes significantly enriched in SKGNur GFP^hi^ T.4N*_Nr4a1_* cells (see fig. 5A) for two replicate mice in each subgroup.

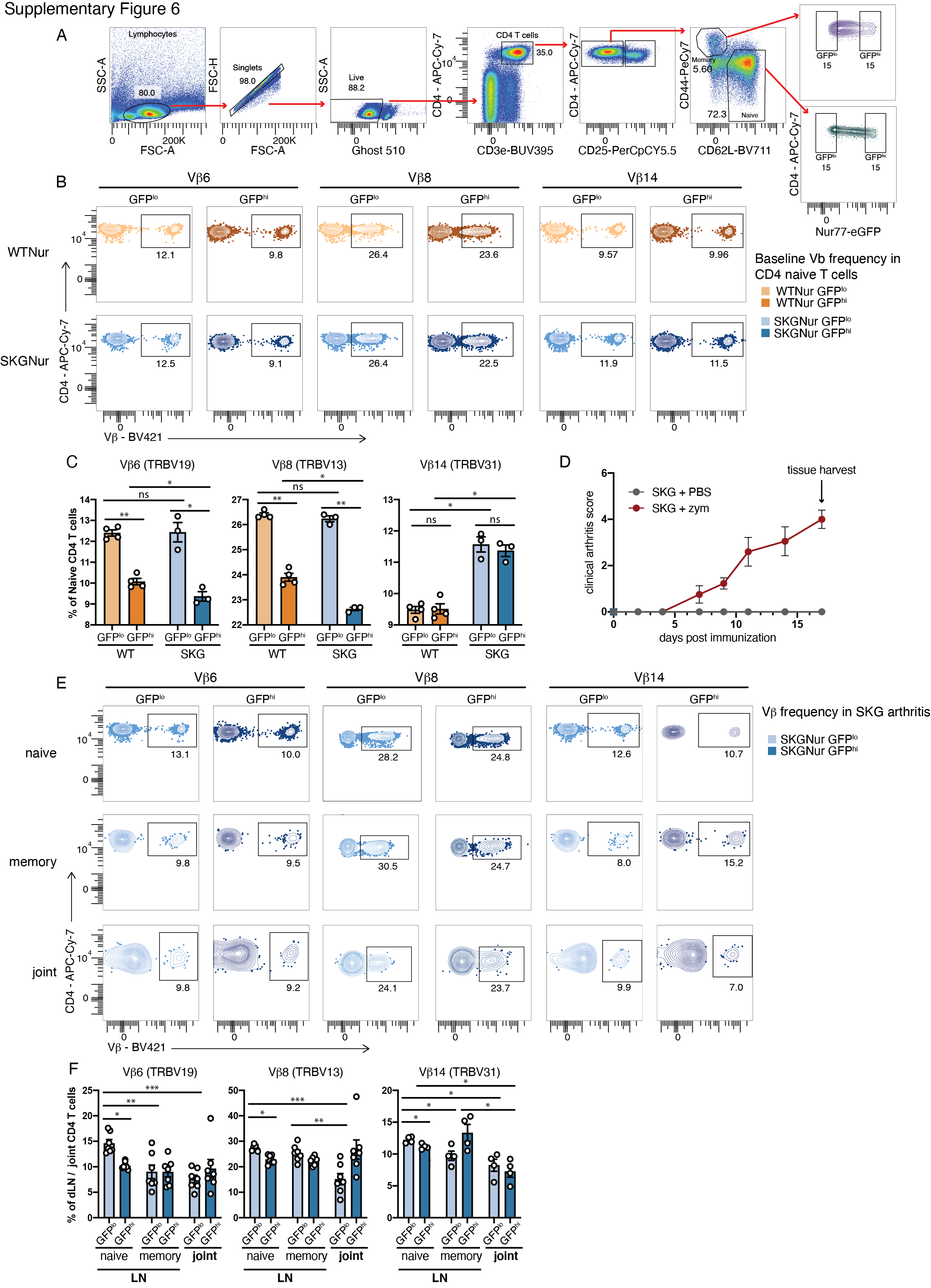

**Supplementary Fig. 6. TCR Vb’s unresponsive to Balb/c MMTV superantigen do not expand in cells marked by TCR signaling reporter.** (**A**) Flow cytometry gating used to identify GFP^hi^ and GFP^lo^ populations in naïve (CD62L^hi^CD44l^o^) and memory (CD44^hi^CD62L^lo^) CD4+CD25- T cells for Vb identification in WTNur and SKGNur lymphocytes. (**B-C**) Representative FACS plots (**B**) of naïve peripheral CD4 T cells with indicated TCR Vb protein usage determined by flow cytometry in GFP^lo^ and GFP^hi^ T cells from LN of WTNur and SKGNur mice prior to arthritis induction and quantified in (**C**) where bar graphs depict mean frequency (± SEM), n = 3-4 mice per group, experiment repeated at least 3 times. (**D**) Arthritis score in SKGNur mice ± i.p. zymosan (red) or PBS (grey), n=4 mice in each group, representative of at least 3 experiments. (**E-F**) Representative FACS plots (**E**) of peripheral naïve or memory, or joint CD4 T cells with indicated TCR Vb protein usage determined by flow cytometry in GFP^lo^ (light blue) and GFP^hi^ (dark blue) T cells from LN or joints of SKGNur mice 2.5 weeks after arthritis induction with zymosan (as seen in Supplemental Fig. 6D) and quantified in (**F**) where bar graphs depict mean frequency (± SEM), n = 7 mice per group pooled from 2 experiments. (C, F) Significance indicated by asterisk for *P* value (exact permutation test for unpaired samples) or FDR (paired t-test) < 0.05 (*), < 0.1 (**), or < 0.001 (***)**.**

Supplementary Data

Data S1. (data_S1_bulk_RNA_seq_diff_exp.xlsx)

Differential expression results for all bulk RNA seq comparison.

Data S2. (data_S2_heatmap_gene_list_with_modules.csv)

List of genes in heatmap in **Fig. 1C**.

Data S3. (data_S3_GSEA_reports.xls)

GSEA reports for all GSEA analyses.

Data S4. (data_S4_scRNAseq_diff_genes_by_cluster.csv)

Differential expression results for analyses in **Fig. 2E-F**.

Data S5. (data_S5_fig_2_diff_gene_lists.xlsx)

Differential expression results for single cell RNA sequencing clusters.

Data S6. (data_S6_diff_exp_Tnfrsf9_pos_Egr2_pos.csv)

Differential expression results for analysis in **Fig. 3C**.

Data S7. (data_S7_top_300_heatmap_gene_list.tsv)

List of genes in heatmap in **Fig. 4F**.

Data S8. (data_S8_gini_coefficients.csv)

Gini coefficients for clonotype distribution for cells in each cluster from each mouse.

Data S9. (data_S9_TRBV_paired_test_nr4a1_cluster.csv)

Statistics for comparisons of TRBV frequencies shown in **Fig. 5A.**

Data S10. (data_s10_diff_exp_SKG_High_v_SKG_Low_for_TRBV.xlsx)

Differential expression results for analysis in **Fig. 5G and Supplementary Fig. 4D.**
